# Mesenchymal Stem Cell-Mediated Intercellular Communication: Mapping the Interactome for Skeletal Muscle Homeostasis and Regeneration

**DOI:** 10.1101/2025.10.02.678862

**Authors:** Xingyu Liu, Edgar E. Perez Carbajal, Yih-Chii Hwang, Benjamin W. Gilman, Sarah A. Bliss, Christapher S. Morrissey, Michael N. Wosczyna

## Abstract

Mesenchymal stem cells (MSCs) support tissue homeostasis and regeneration, yet their molecular signals remain largely enigmatic. In skeletal muscle (SkM), MSCs, known as fibroadipogenic progenitors (FAPs), are essential for maintenance and repair, orchestrating these processes through intricate cellular communication networks. Given the critical role of SkM in lifelong health and longevity, FAP signaling has drawn significant interest as a potential therapeutic target and a model for MSC interactions. However, deciphering FAP-derived regulatory signals remains challenging due to their pleiotropic complexity. Here, we employ a systems-based approach to construct a comprehensive FAP interactome in both homeostatic and regenerating SkM. By integrating unique single-cell RNA sequencing atlases with advanced computational analyses, we identify putative FAP-mediated signaling pathways and validate their biological relevance through FAP depletion experiments, assessing disruptions in key pathways. This approach reveals novel signaling networks across diverse SkM cell populations, corroborates key FAP interactions from recent studies, and provides a valuable dataset for modeling MSC interactions and their roles in SkM homeostasis and regeneration.

## Introduction

Skeletal muscle (SkM) is one of the largest organs in the human body, providing force for locomotion, glycogen storage for energy reserves, and involuntary movement for thermoregulation^1–3^. The robust activities of tissue stem cells enable SkM to maintain and regenerate its structure, allowing these functions to be preserved following injury^4^. However, these maintenance and regenerative capabilities diminish with age and pathology, negatively impacting SkM function^5–9^.

At the cellular level, SkM maintenance and regeneration rely on satellite cells (SCs, also known as muscle stem cells) for the continued supply of progenitors that differentiate into myoblasts to maintain or rebuild myofibers following injury^10^. Recently, mesenchymal stem cells (MSCs) located in the stromal space of SkM, known as fibroadipogenic progenitors (FAPs), have been proven as active participants in these processes, essential for SkM maintenance and regeneration^11,12^. These findings were made possible by using the platelet-derived growth factor receptor alpha (PDGFRα) gene locus to specifically target FAPs in vivo for conditional labeling and ablation^12,13^. When FAPs are depleted, muscle atrophies and SC numbers decrease over time, indicating that FAPs are required to maintain SkM and the SC niche in homeostatic conditions^12^. Likewise, injury of FAP-depleted SkM reveals delayed regeneration, with the underlying regenerative cellular milieu exposing limited expansion of SCs and decreased infiltration of immune cells^12^. These data support the requirement of FAPs in SkM maintenance and regeneration, presumably through cell-extrinsic signaling. While the necessity of FAPs has been soundly established for SkM health, the in vivo molecular pathways for which FAPs modulate the homeostatic and regenerative cellular milieus remain incomplete.

Emerging data suggest that a multitude of molecular mechanisms are modulated by FAPs, putatively establishing these cells as master regulators of SkM health. FAPs have been identified as key producers of the extracellular matrix (ECM), which supports the self-renewal and maintenance of SCs during muscle regeneration^14–16^. However, while ECM production facilitates the regenerative process, excessive ECM deposition is associated with fibrosis in pathological states^17,18^. Analyses by single-cell RNA-sequencing (scRNA-seq) of cells in SkM confirmed the expression of ECM genes by FAPs during homeostasis and a subsequent upregulation during regeneration^19^. The proliferative expansion of FAPs during the early days of regeneration following injury is essential for transforming growth factor beta (TGF-β)-dependent ECM deposition, but FAP numbers must be tightly controlled by infiltrating macrophages that secrete tumor necrosis factor to induce FAP apoptosis^20^. Furthermore, FAP secretion of IL-6 and WISP1 has been demonstrated to increase SC proliferation and differentiation^11,21,22^. In homeostatic muscle, a subpopulation of DPPIV+ FAPs sustains colony-stimulating factor 1 (CSF1) expression in the local niche, supporting the self-renewal of tissue-resident macrophages^23^. Interestingly, in aging scenarios, p16+ FAPs may recruit macrophages through the secretion of C–C chemokine ligand 2 (CCL2) and osteopontin and prime their polarization towards the M2 phenotype^24^. These data have exemplified the importance of FAP communication with cells of the homeostatic and regenerative milieu.

While FAPs are essential for SkM health and many potential cell-to-cell signaling pathways have been putatively uncovered, it remains challenging to effectively decipher the FAP interactome due to the complexity of in vivo homeostatic and regenerative environments coupled with the pleiotropy of FAP-dependent signals. However, the capacity to interpret multicellular interactions has substantially increased with the advancement of computational programs capable of analyzing cell intrinsic transcriptomes and predicting cell-to-cell communication based on research-informed signaling networks.

In this study, we generate transcriptome atlases for all mononuclear cells in the SkM homeostatic and regenerative milieu with FAPs present, and with FAPs depleted using our in vivo targeting approach. These atlases provide datasets to infer interactions and molecular mechanisms and then determine if FAP depletion functionally alters these presumed communications. By incorporating the FAP-depleted atlases, we provide a means to establish a multicellular and biologically relevant FAP-dependent interactome for mononuclear cells in homeostatic and regenerative milieus. Our data uncover remarkable intercellular communication pathways between FAPs and other cells of the milieu, namely, SCs, macrophages, neutrophils, and antigen-presenting cells (APCs). These pathways were elucidated through the increased resolution of cellular subpopulations in scRNA-seq modeling and the capacity of our FAP-depleted datasets to confirm the reliance of the predicted mechanism on FAP presence in an all-inclusive cellular assessment in SkM. Furthermore, our datasets provide a resource for the SkM field to predict, test, and validate FAP signaling in homeostatic and regenerative environments. In the following sections, we will present the generation of these datasets and the elaborate bioinformatics approach used to confirm and reveal new FAP mechanisms regulating the SkM cellular milieus.

## Results

### In vivo targeting of FAPs for depletion

Given that PDGFRα expression is highly specific to MSCs, we utilized our previously reported PDGFRα^CreER^ mice, where the translation of CreER is driven by the endogenous start codon in the PDGFRα locus, to target FAPs in SkM for depletion. We crossed the PDGFRα^CreER^ mice with R26^DTA^ mice to produce PDGFRα^CreER^; R26^DTA^ offspring, referred to, hereafter, as PACE;DTA mice (Fig. 1A). We previously demonstrated that following tamoxifen administration to PACE;DTA mice, FAP depletion was highly efficient^12^. We confirmed this result and demonstrated an ∼85% depletion of SkM FAPs (Fig. 1B). Henceforth, tamoxifen-treated PACE;DTA will be referred to as FAP-depleted (in text) or “Depleted” (in figures) mice.

**Figure 1.**
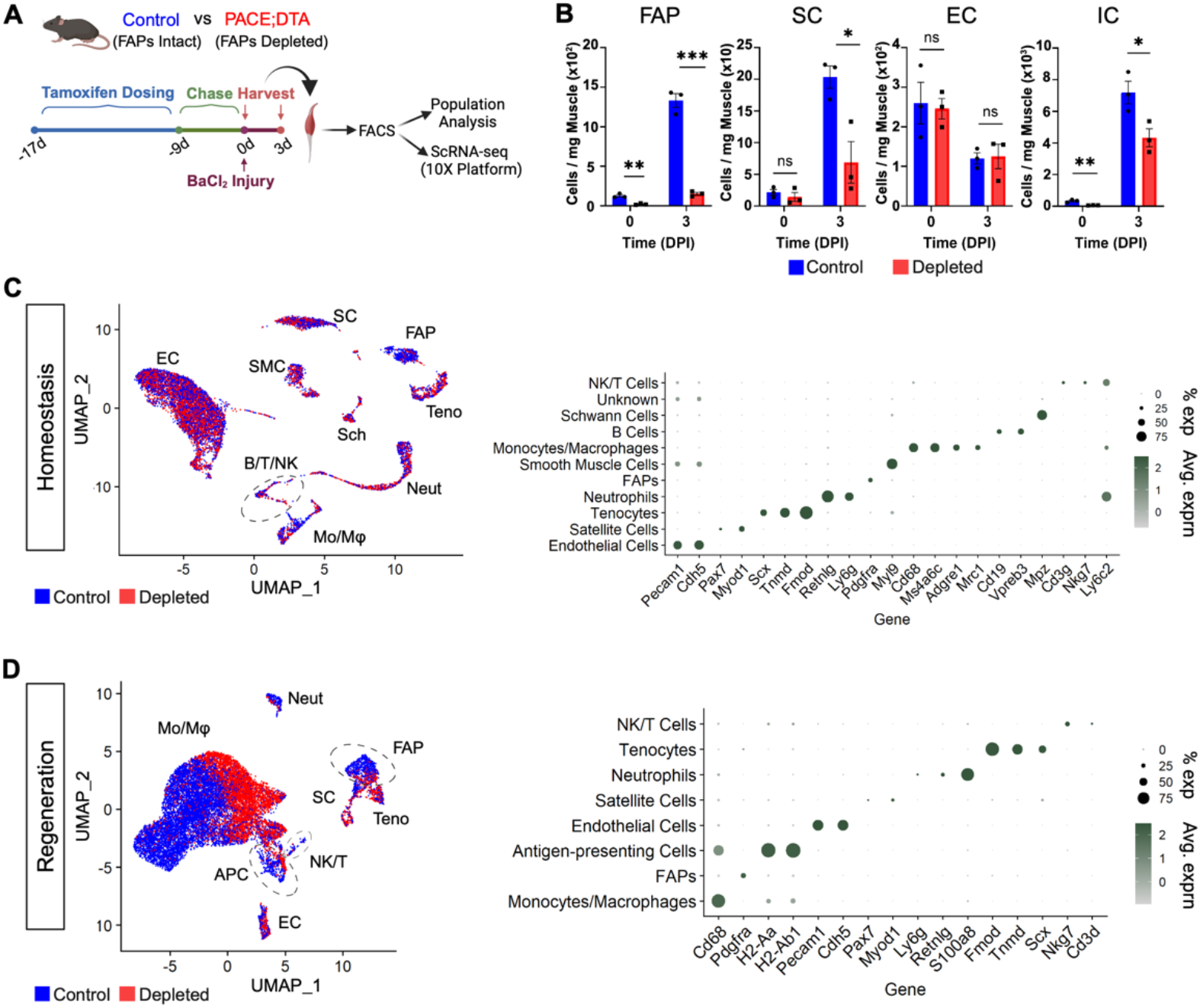
Developing transcriptional atlases to interrogate the FAP interactome in SkM homeostasis and regeneration. (A) An overview of the experimental scheme. Note, cells were harvested from uninjured and 3 days post injury (dpi) SkM. (B) Quantitation by FACS of noted cell types from FAP containing (Control) and depleted mice. n=3 per condition. (C, D) 10X scRNA-seq data of SkM cell populations represented in UMAPs with dot plots quantifying the marker genes used for cluster identification in (C) homeostasis and (D) regeneration. Single cells are color-coded for control (blue) or FAP-depleted (red) origin. Each condition has cells combined from an n=2. *p ≤ 0.05, ** p < 0.01, ***p < 0.001. Error bars represent SEM.

### Cellular response to FAP depletion in homeostatic and regenerating SkM

We next examined the cellular profile of uninjured and injured (3 days post-injury (dpi)) SkM from FAP-depleted and control mice with fluorescence-activated cell sorting (FACS) (Fig. 1A, B). All cohorts of mice received tamoxifen to control for any related toxicity. For injuries, BaCl_2_ was injected intramuscularly following the tamoxifen chase period to avoid the possibility of de novo recombination and DTA-directed cell death during the injury and in the regenerative period. The uninjured and injured lower hindlimb muscles were collected and processed for mononuclear cell extraction. In FAP-depleted muscle, SC numbers were unaffected before injury but exhibited significantly impaired expansion upon injury with a 66% deficit when compared to control at 3 dpi (Fig. 1B). Under homeostatic conditions, only small numbers of hematopoietic cells labeled with the pan-immune marker CD45 were present in muscle, and these numbers further diminished following FAP depletion. Furthermore, after injury, the robust immune cell response exhibited significantly impaired expansion in FAP-depleted mice with a 40% deficit at 3dpi when compared to control. FAP depletion, regardless of injury status, did not affect the number of endothelial cells (Fig. 1B). These cellular responses following FAP depletion are consistent with findings from previous reports^12,13^.

### Establishing single-cell FAP-dependent transcriptome atlases for uninjured and injured SkM

Live cells were collected by FACS from SkM of FAP-depleted and control mice before and after injury (3 dpi) for cellular profiling. Using the Chromium 10X platform, we performed scRNA-seq on all cells from each experimental cohort. Standard processing was conducted with the Seurat R package^25^, integrating scRNA-seq data from two biological replicates per sample. After quality control, we retained 6,145 control and 6,004 FAP-depleted cells from uninjured samples, and 10,294 control and 10,120 FAP-depleted cells from 3 dpi samples for analysis. As defined above by FACS profiling, endothelial cell numbers remain unchanged between FAP-depleted and control groups at 3 dpi (Fig. 1B). Therefore, the depleted scRNA-seq dataset in the injured condition was normalized to match the control, maintaining the same endothelial cell count so that the observed lack of expansion when FAPs are depleted was accurately reflected in the scRNA-seq modeling. The Uniform Manifold Approximation and Projection (UMAP) method was used for dimensional reduction, and the Louvain algorithm for community detection, processing uninjured and injured samples independently. We identified ten distinct populations in uninjured samples and eight in injured samples (Fig. 1C, D; Supplementary Fig. 2A, B).

## FAPS IN HOMEOSTATIC SkM

### FAP depletion in homeostasis induces a phenotype switch of SkM resident immune cells

We analyzed the size of each cell population in SkM from FAP-depleted and control mice. An 81% reduction of FAPs was observed in FAP-depleted mice, consistent with our flow cytometry results, and this depletion impacted all FAP subpopulations equally (Supplementary Fig. 3A). SC quantities remained consistent between groups, most likely due to this being a short-term depletion versus our previous report of a cumulative reduction in SCs when FAPs were depleted for 9 months^12^. Interestingly, FAP depletion resulted in a redistribution of the immune cell population, with a 43% and 24% decrease in CD68+ monocytes/macrophages (Mo/Mφs) and CD19^+^ B cells, respectively. The number of CD3+ NK/T cells and Schwann cells were increased by 43% and 38%. No appreciable changes in population size were observed in other cell types (Fig. 2A, B).

**Figure 2.**
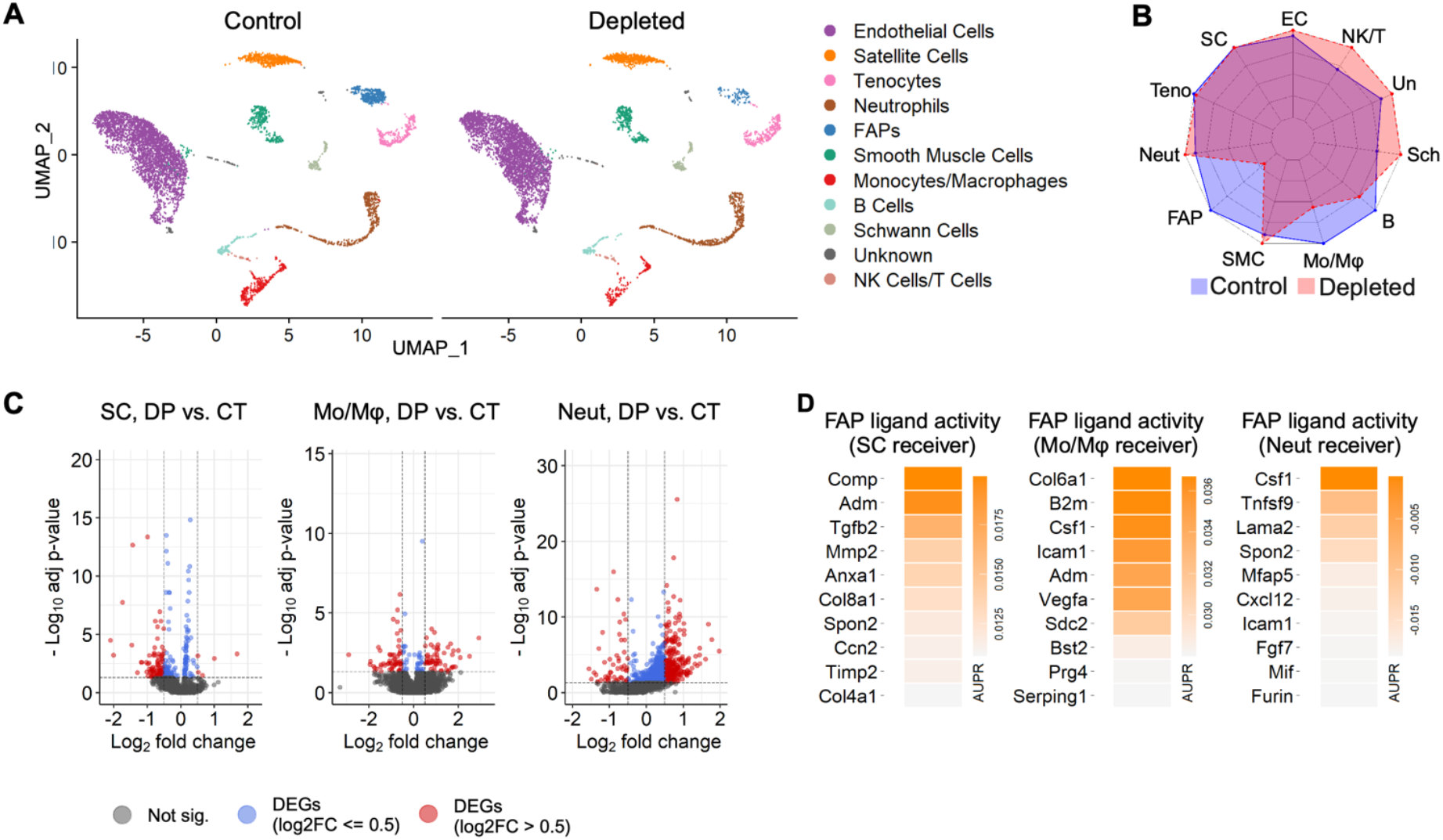
Deciphering the impact of FAPs on cellular populations in homeostatic SkM. (A) UMAPs of cell populations from uninjured SkM with FAPs present (control) or depleted. (B) Radar plot comparing population frequencies in control vs. FAP-depleted SkM. (C) Volcano plots representing DEGs in the noted cell populations from control and FAP-depleted samples. Red: DEGs with log_2_FC > 0.5 and adjusted p-value < 0.05. Blue: DEGs with log_2_FC ≤ 0.5 and adjusted p-value < 0.05. Gray: DEGs with adjusted p-value ≥ 0.05. (D) Top 10 FAP-derived ligand activity scores in uninjured SkM quantitated by the area under the precision-recall curve (AUPR), with noted cell populations set as the receiver.

### FAP depletion in homeostasis alters the gene expression in SCs, Mo/Mφ, and neutrophils

Next, we conducted differentially expressed gene (DEG) analyses on each population in SkM to investigate the potential intracellular signaling alterations caused by FAP depletion. SC, Mo/Mφs, and neutrophils underwent significant transcriptomic profile changes, while others remained unaffected (Fig. 2C and Supplementary Fig. 3B). Functional enrichment analyses using these DEGs demonstrated significant (adj. p-value < 0.05) biological pathways enriched in all three populations when FAPs were depleted (Supplementary Fig. 3C-E). Interestingly, collagen-associated pathways were upregulated in SCs. At the same time, adhesion and maintenance of location were downregulated, suggesting SCs are preparing to exit the niche, consistent with previous research that indicates a loss of SCs over time with FAPs depleted. Mo/Mφs in FAP-depleted muscle showed comprehensive enrichment in chemotaxis-associated activities, indicating potential compensatory leukocyte recruitment for microenvironment and niche maintenance. However, pathways involved in the immune defense process were down-regulated in the Mo/Mφs from the FAP-depleted dataset, suggesting an impaired capacity for immune surveillance and clearance of apoptotic cells in muscle lacking FAPs (Supplementary Fig. 3D). Neutrophils from the FAP-depleted muscle were enriched for cell cycle activation but with weak migration (Supplementary Fig. 3E), which suggests neutrophils tend to remain in an immature state when FAPs are absent.

### Elucidating the interactome between FAPs and SCs, and FAPs and Mo/Mφs

#### FAPs and SCs

Previous research by our group reported a significant decline in SCs 9 months after FAP depletion^12^. This finding led us to consider the role of FAPs in maintaining a beneficial niche for SCs. As mentioned, our single-cell level analysis did not reveal significant changes in the number of SCs shortly following FAP depletion but did resolve a change in ECM and adhesion characteristics (Fig. 2B and Supplementary Fig. 3C), consistent with our FACS results (Fig. 1B), suggesting the impact of the loss of FAPs on SCs is cumulative over time. We then hypothesized that the cues originating from FAPs to maintain SCs are continually needed and can be revealed by examining gene expression changes intrinsic to SCs. Therefore, we employed the computational tool NicheNet^26^ to investigate the effects of active ligands from FAPs on SCs. NicheNet scores and ranks ligands from sender cells, in this case FAPs, identifying sender-derived signals of the highest potential to regulate DEGs in receiver cells from alternative conditions, here, DEGs of SCs from FAP-depleted versus control muscle. In addition to SCs, we also identified ligands from FAPs which may regulate the transcriptomes in Mo/Mφs and neutrophils (Fig. 2D). While most ligands for SCs and Mo/Mφs have positive values for area under the precision-recall curve (AUPR), those for neutrophils are primarily negative, meaning FAP-derived ligands are likely not directly responsible for the altered transcriptome in neutrophils from depleted mice. As such, we focused on FAP communication with SCs and Mo/Mφs in homeostatic SkM.

The top FAP ligands (Mmp2, Col8a1, Spon2, Ccn2, Timp2) predicted in our NicheNet analyses to influence SCs are also expressed by other cell types in SkM, notably tenocytes (Fig. 3A). These data suggest that alternative cellular sources cannot compensate for FAPs in the production of ligands needed to maintain SCs over time, and in the FAP-depleted background the cumulative impact on the stem cell niche leads to a loss of SCs. To further investigate FAP communication with SCs, we implemented the computational tool, CellChat^27^, an inference method that does not require the cell-intrinsic analyses of receiver cells as input, but instead takes advantage of the expression levels of ligand-receptor pairs. This approach provides insight into possible cell-cell communication, though it does not necessarily indicate actual interactions or functional impacts on the receivers. CellChat analyses using control data as input suggested that Type I collagen, primarily encoded by Col1a2 and Col1a1 and secreted by FAPs, contributes to SC maintenance by interacting with receptors Sdc4 and Cd44 on SCs (Fig. 3B), again implying that FAP-dependent ECM contribution is essential in maintaining the niche for SCs to remain in the sublaminar position.

**Figure 3.**
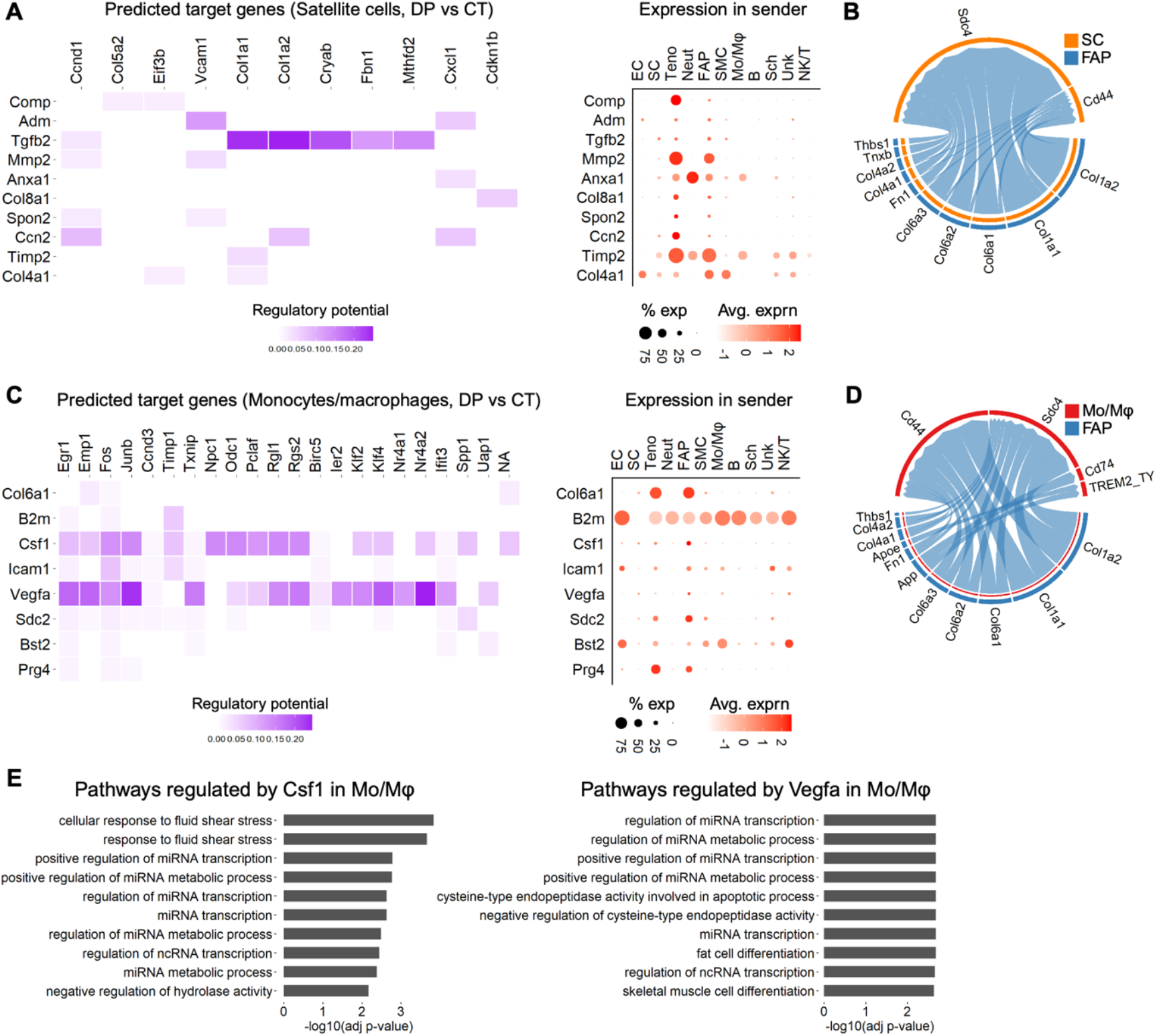
FAP regulatory mechanisms of SC and Mo/Mφ activity in SkM homeostasis. (A, C) Top FAP-derived ligands with the highest activity and their predicted impact on DEGs in (A) SCs or (C) Mo/Mφs, with the dot plots showing the abundance and expression level of the corresponding ligands across the cells in uninjured SkM. (B, D) Interactions between FAPs (sender) and (B) SCs (receiver) or (D) Mo/Mφs (receiver). Line thickness represents interaction strength. (E) Top 10 biological processes in Mo/Mφs most likely to be regulated by Csf1 (left) and Vegfa (right). Truncated terms “TREM2_TY” = TREM2_TYROBP, and “cysteine-type endopeptidase activity involved in apoptotic process” = negative regulation of cysteine-type endopeptidase activity involved in apoptotic process.

#### FAPs and Mo/Mφs

For FAP communication with Mo/Mφs in homeostatic conditions, our NicheNet analyses revealed that Csf1 and Vegfa from FAPs have high regulatory potential on the Mo/Mφ population (Fig.3C). Functional enrichment analyses using genes in Mo/Mφs predicted to be regulated by Csf1 and Vegfa showed that both induced pathways were associated with microRNA (miRNA) transcription and processing specific to monocytes and macrophages (Fig. 3E). While miRNAs were not detected in our differential expression analyses, this is likely due to the limitation of 10X sequencing to effectively capture small RNAs. In addition, CellChat analyses revealed that FAP-derived collagen signaling also has an active role in FAP-Mo/Mφ communication, in which Col1a1, Col1a2, Sdc4, and Cd44 are key contributors (Fig.3D). These data suggest that FAPs, and tenocytes, engage most actively with other cell populations, primarily in an ECM-receptor manner (Supplementary Fig. 4A, B).

### Resolving subpopulations of SCs, Mo/Mφs, and neutrophils impacted by FAP depletion

#### FAPs and SC subpopulations

Since heterogeneity in cell phenotype within a target population may provide finer insight into the role of FAP signaling in homeostasis, we independently reclustered SCs, Mo/Mφs, and neutrophils to explore granularity in each population. For SCs, 5 clusters were resolved, with a single cluster (noted as cluster 2 in Supplementary Fig. 5A, B) showing an appreciable decrease in cell number when FAPs were depleted (Supplementary Fig. 5A). Pathway analyses examining the DEGs in cluster 2 were mainly centered around protein processing (Supplementary Fig. 5C). Two additionally enriched pathways were muscle organ and connective tissue development. Examination of myogenic genes in this population leads us to hypothesize these cells are in a transient state between quiescence and poised to activate (still quiescent) (Supplementary Fig. 5D). These data suggest that FAP depletion leads to the preferential loss of this SC subpopulation in homeostasis that is responsive to ECM levels and responsible for muscle maintenance.

#### FAPs and Mo/Mφ subpopulations

For Mo/Mφs, we observed 6 subpopulations following reclustering. Interestingly, 3 subpopulations of macrophages declined in FAP-depleted muscle, with two of these being responsive to interferon beta (IFN-β) and interferon gamma (IFN-γ), indicating that FAPs are required to maintain macrophages in SkM. Conversely, infiltrating monocytes increased in FAP-depleted samples, possibly indicating a compensatory mechanism attempting to repopulate tissue macrophages (Supplementary Fig. 5E, F).

Functional enrichment analyses further confirmed the identities of subpopulations (Supplementary Fig. 5G). Note, that the “Mφ” population lacked enriched pathways, but shared many expressed marker genes with IFN-β responsive macrophages (Supplementary Fig. 5F), potentially indicating a common origin for these two subpopulations. Collectively, these data implicate FAPs in maintaining resident Mo/Mφs, a finding made possible by the increased resolution of cell populations by scRNA-seq, but more importantly, emphasizing the incorporation of our FAP-depleted atlas to test and verify predicted interactions beyond just typical transcriptional correlation.

#### FAPs and neutrophil subpopulations

Neutrophil reclustering resolved 4 subclusters, each representing different stages of neutrophil development (Supplementary Fig. 6A, B). To define their progression through maturation, using Monocle 3^28^, we applied an alternative method of dimensional reduction, performed trajectory inference on the resulting low-dimensional data, and calculated the pseudotime value for each cell with immature neutrophils designated as the origin (Supplementary Fig. 6C, D). Interestingly, proliferating neutrophils followed a distinct trajectory separate from the main trajectory (Supplementary Fig. 6C, D), suggesting that these neutrophils may originate from different parent populations. The expression of immature neutrophil marker genes gradually decreased over pseudotime, whereas the expression of mature neutrophil marker genes progressively increased (Supplementary Fig. 6E), confirming the accuracy of the cell ordering. Moreover, the pseudotime values of neutrophils and pathway enrichment analysis correspond to their assigned identity (Supplementary Fig. 6F, G). Although the total number of neutrophils is similar between the FAP-depleted and control muscle, the control dataset contains a higher proportion of mature neutrophils (Ccl6, Cxcl2, Il1b; Supplementary Fig. 6A, B, E), while the FAP-depleted dataset has more immature neutrophils (Ngp, Camp, Ltf; Supplementary Fig. 6A, B, E). Collectively, our data underscore a potential role for FAP signaling regulating neutrophil development in homeostatic SkM.

Though FAP-expressing signals are not considered to make a significant impact on the neutrophil transcriptome in the short-term, communication through FAP expression of Cxcl1 ligand and its receptor Cxcr2 expressed by neutrophils was clearly defined by CellChat analyses (Supplementary Fig. 6H). Meanwhile, the capacity of leukocytes to generate chemokines is substantial^29^ and thus, inter-leukocyte communication is highly likely. Therefore, we explored the potential communication pathways between macrophages and neutrophils. These analyses revealed that in control conditions (FAP-containing), Mo/Mφs express chemokine ligands Ccl6, Ccl9, Cxcl2, and Ccl7, which mainly target Ccr1 and Cxcr2 expressed by neutrophils (Supplementary Fig. 6I). Interestingly, the expression of these ligands is lower in Mo/Mφ from FAP-depleted mice, highlighting the potential of a secondary, Mo/Mφ-dependent impact on neutrophils (Supplementary Fig. 6I). We also found that neutrophils express Ccl6, which can recruit Mo/Mφs by their expression of the receptors Ccr2 and Ccr1 (Supplementary Fig. 6J). Similarly, the number of Ccl6 transcripts also declined in the neutrophils in FAP-depleted mice, which is consistent with our observation that the number of mature neutrophils (marked by Ccl6 expression) was lower in FAP-depleted mice (Supplementary Fig. 6J). Our findings suggest that FAPs contribute to neutrophil recruitment both directly, albeit minimally, and indirectly, while also playing a role in their maturation within homeostatic SkM, helping to maintain a balanced and effective immune environment. Of note, previous studies have only reported low numbers of neutrophils in uninjured tissue, whereas our data detects a palpable presence of these cells in homeostatic SkM. Our study did not include perfusion prior to isolating SkM in an effort to limit any technical impact of this procedure on the tissue and keep processing as rapid as possible to maintain transcriptional profiles in cells. Thus, the neutrophil population may also contain some cells from circulation.

## FAPS IN REGENERATING SkM

### FAPs are crucial for the regenerative response of SCs and many immune cell populations

Similar to our approach above for homeostasis, we first examined our scRNA-seq data from regenerating muscle (3 dpi) for the cellular response of entire populations to FAP depletion. We confirmed FAPs remained depleted during regeneration with a 69% reduction that cannot be rescued by the remaining FAPs through proliferation, consistent with previous reports^12^. SC numbers were reduced by 21%, and the reduction in immune cells noted by FACS was further resolved in this scRNA-seq dataset, demonstrating a decrease in Mo/Mφs, APCs, lymphoid cells (mainly NK and T cells), and neutrophils by 39%, 27%, 78%, and 69%, respectively. Interestingly, tenocytes were the only cell population demonstrating a positive response to FAP depletion, increasing by 46% (Fig. 4A, B).

**Figure 4.**
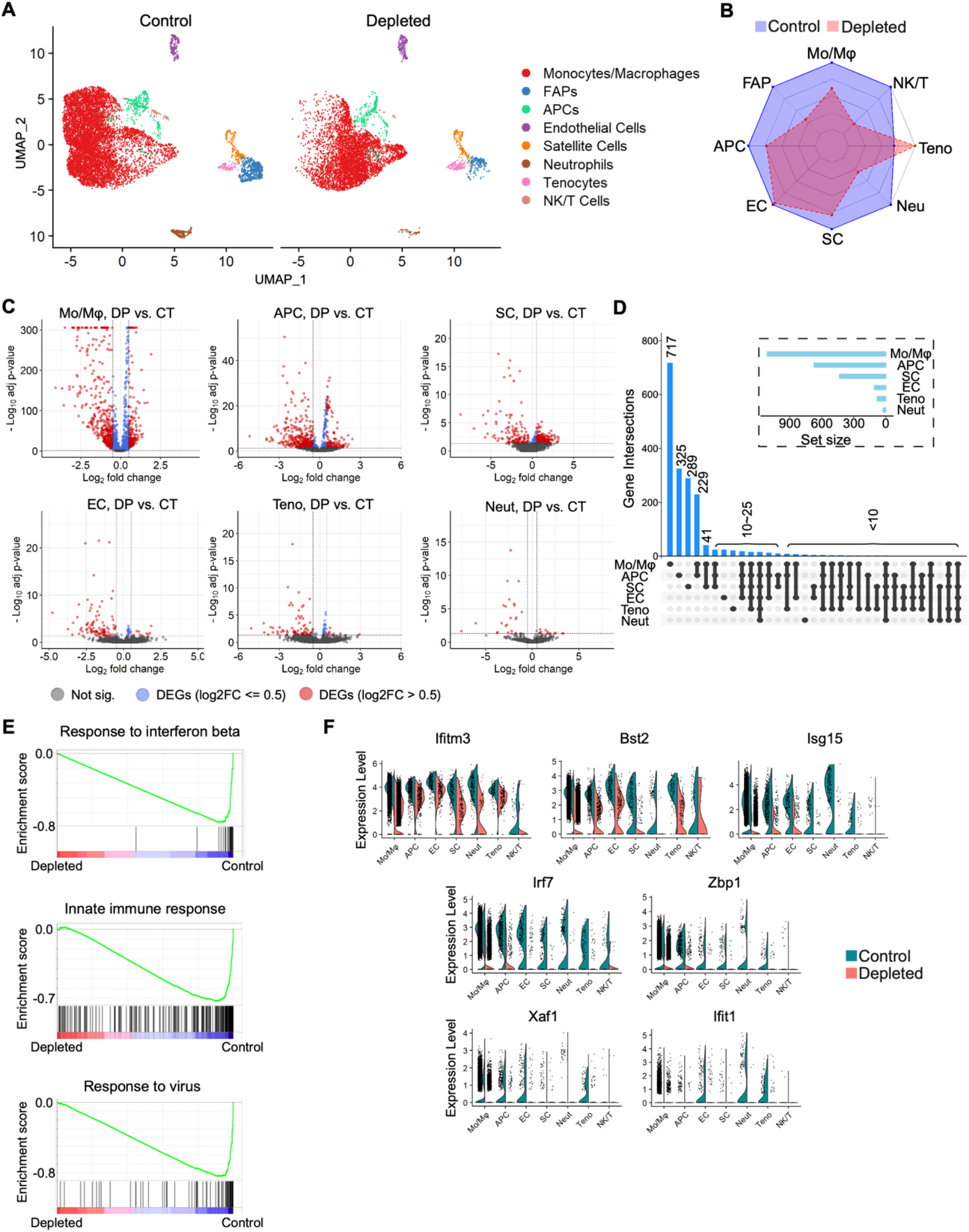
Resolving the FAP-dependent cellular regulation in SkM regeneration. (A) UMAPs derived from scRNA-seq of cells from FAP-containing (control) or depleted SkM at 3 dpi. Cell colors correspond to their noted identity. (B) Radar plot showing the relative difference in the number for each cell population noted in “A.” (C) Volcano plots reveal DEGs in the noted cell populations by comparing cells from depleted SkM to their control counterparts. Red: DEGs with log_2_FC > 0.5 and adjusted p-value < 0.05; Blue: DEGs with log_2_FC ≤ 0.5 and adjusted p-value < 0.05; Gray: DEGs with adjusted p-value ≥ 0.05 (D) Unique DEGs shared among specific cell populations. The main plot illustrates the number of unique DEGs shared exclusively among the cell populations indicated by black circles, with shared DEGs indicated by connected circles. The inset represents the total number of DEGs identified in each cell population. (E) GSEA plots demonstrate the downregulation of immune-related biological pathways in FAP-depleted SkM at 3 dpi compared to control counterparts. (F) Expression levels of IFN-β response-associated genes across individual cell populations, split by control and depleted samples.

### FAPs depletion in regeneration alters the gene expression in SCs, Mo/Mφs, and neutrophils

To assess the impact of FAP-depletion on cell-intrinsic mechanisms of cell populations in the regenerative milieu, we examined DEGs in cell populations between FAP-depleted and control mice. FAP depletion resulted in significant changes in SCs, Mo/Mφs, and APCs, while tenocytes, endothelial cells, and neutrophils were only moderately impacted (Fig. 4C, D). The remarkable overlap in DEGs between Mo/Mφs and APCs suggests they have shared regulatory pathways, putatively reflecting a close relationship in their development or response to stimuli (Fig. 4D). Although lymphocytes exhibited a reduction in size, no notable changes in gene expression were observed, suggesting a deficiency in their replenishment from external sources, rather than alterations in proliferation processes.

### FAP depletion leads to a global reduction of immune response pathways

In the context of regeneration, we detected a striking down-regulation of immune response-associated pathways in cells of FAP-depleted muscle when compared to those in control muscle (Fig. 4E and Supplementary Fig. 7A-F). In particular, the IFN-β-stimulated transcriptomic program was attenuated across all cell populations within the FAP-depleted muscle. Furthermore, genes involved in the cellular response to type I IFN exhibited lower expression in the FAP-depleted mice irrespective of cell type (Fig. 4F). These results demonstrate that FAPs are indispensable for the activation of cellular populations in muscle through IFN-β stimulation following injury.

### Elucidating the interactome between FAPs and SCs, tenocytes, endothelial cells, Mo/Mφs, and APCs

To explore the potential for FAP-specific ligands to interact with target cells and alter their gene expression, we again utilized the computational tool, NicheNet. Of note, the Mo/Mφ and the APC clusters both express high levels of CD68, potentially indicating a shared progenitor origin. (Fig. 1D). Thus, they are collectively henceforth termed “mononuclear phagocytes”^30,31^. Our analyses revealed that Il6 is near uniquely expressed in FAPs and has a significant potential to regulate a substantial number of DEGs in almost all cell types in regenerating SkM (Fig. 5A, D, Supplementary Fig. 8A, Supplementary Fig. 11A, D, and Supplementary Fig. 12A). Fgf7 expression is also specific to FAPs, but it was only predicted to impact non-blood cells (SCs, tenocytes, and endothelial cells). The receptors for Il6 and Fgf7 (Fgfr1, Fgfr4, Nrp1, Il6ra, Il6st) are highly expressed across these cell populations (Supplementary Fig. 8B). While both Fgf7 and Il6 are involved in immune defense programs, particularly IFN-β- and IFN-γ-stimulated responses, Fgf7 is primarily associated with antiviral responses, and Il6 is linked to pathogen processing and presentation (Fig. 5B, Supplementary Fig. 11C, F, and Supplementary Fig. 12C). We also performed CellChat analysis to infer potential FAP interactions further, using control data as input. As in homeostasis, collagens from FAPs are indicated to influence all cell populations in the regenerative environment (Fig. 5C, Supplementary Fig. 9A, B, Supplementary Fig. 11B, E, and Supplementary Fig. 12B). FAP-derived Fn1 and Thbs2 were also predicted to target several different cell types with the exception of endothelial cells. These data again emphasize the essential pleiotropic signaling originating from FAPs that coordinate the regenerative response.

**Figure 5.**
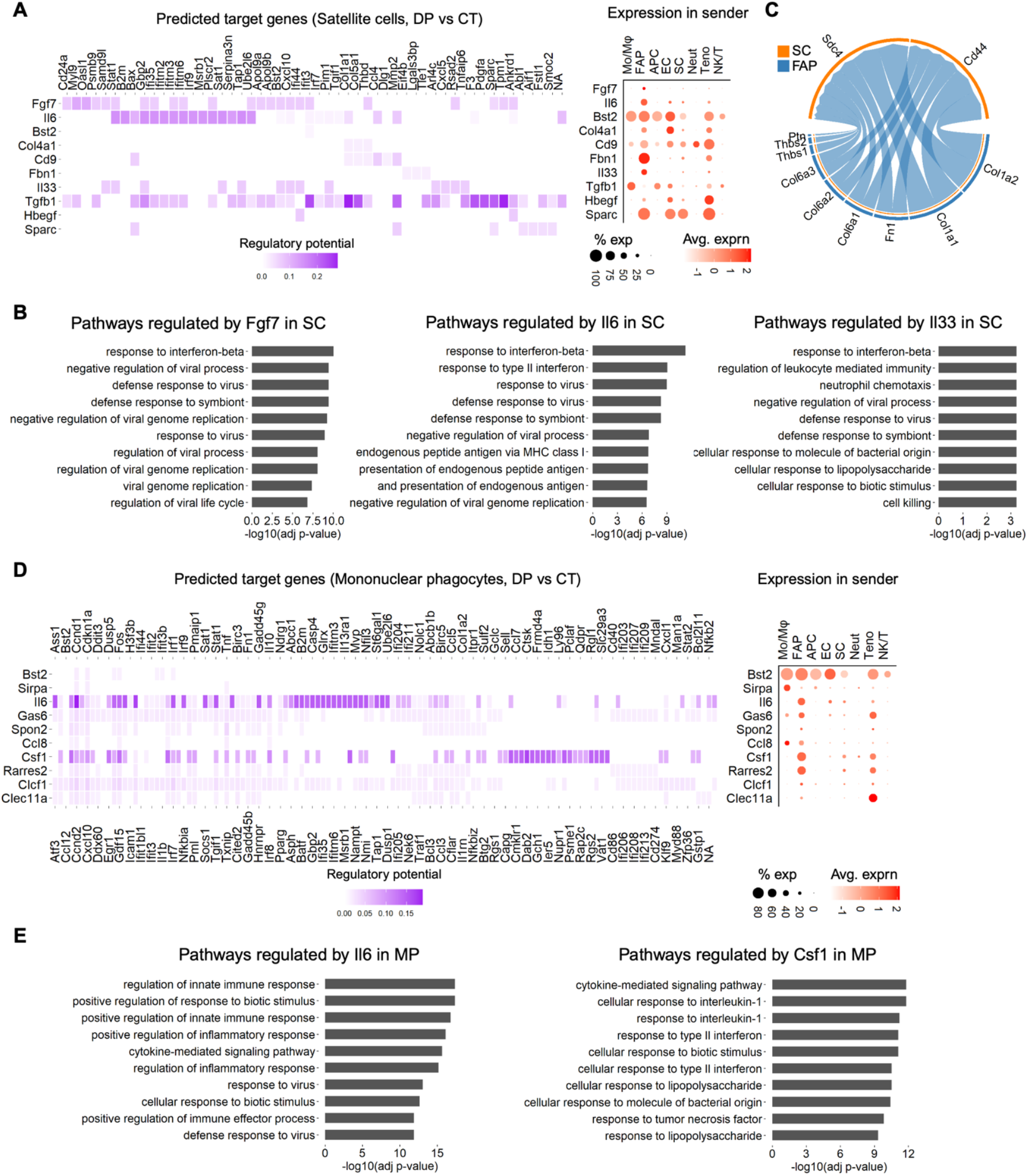
FAP-dependent regulation of SCs and mononuclear phagocytes in injured SkM. (A) Top FAP-derived ligands with the highest activity and their predicted impact on DEGs in SCs, with the dot plot showing the abundance and expression level of these ligands across cell types in SkM at 3 dpi. (B) Top 10 biological processes in SCs inferred to be regulated by Fgf7 (left), Il6 (middle), and Il33 (right). (C) Interactions between FAPs (sender) and SCs (receiver). The thickness of lines represents interaction strength. (D) Top FAP-derived ligands with the highest activity and their predicted impact on DEGs in mononuclear phagocytes, with the dot plot showing the abundance and expression level of these ligands across cell types in SkM at 3 dpi. “Mononuclear phagocytes” refer collectively to monocytes, macrophages, and APCs. (E) Top 10 biological processes in mononuclear phagocytes inferred to be regulated by Il6 (left) and Csf1 (right). Truncated terms “endogenous peptide antigen via MHC class I” = antigen processing and presentation of endogenous peptide antigen via MHC class I, “presentation of endogenous peptide antigen” = antigen processing and presentation of endogenous peptide antigen, and “and presentation of endogenous antigen” = “antigen processing and presentation of endogenous antigen.”

#### FAPs and SCs

In addition to ligands that can target most cells in the regenerative environment, some signals were shown to be cell-type specific. We observed that Il33 and Fbn1 are uniquely expressed in FAPs and included in the top 10 ligands of the FAP-SC regulatory network. Pathway analysis revealed that Il33 shares a similar function with Fgf7 and Il6, particularly in modulating IFN-β-associated responses. While Fbn1 demonstrated high expression in FAPs, the number of DEGs in SCs predicted to be regulated by this gene was relatively low (Fig. 5A). Consequently, the influence of Fbn1 may not directly alter the cellular processes in SCs. Furthermore, Tgfb1 has a strong regulatory impact on gene expression in SCs (Fig. 5A). However, Tgfb1 was broadly expressed across multiple cell types rather than being restricted to FAPs. Thus, its effect on SCs is likely mediated by numerous cells in the niche, like Mo/Mφ, APCs, and other cell populations. Moreover, Tgfb1 was not predicted to originate from FAPs and directly target Mo/Mφs and APCs, and there was no difference in expression of Tgfb1 in Mo/Mφs and APCs between FAP-depleted and control samples. Thus, an indirect impact of FAPs on Tgfb1 from other cell types in the niche seems implausible.

#### FAPs and mononuclear phagocytes

We also uncovered highly specific expression of Csf1 in FAPs. NicheNet and CellChat analyses both find Csf1 originates from FAPs and targets mononuclear phagocytes in healing SkM (Fig. 5D, E, and Supplementary Fig. 12G, H). Csf1 was previously reported to have a crucial role in maintaining the self-renewal ability of macrophages in homeostatic SkM^23^, but its role in SkM regeneration is not fully understood. Here, we noted that in regenerating SkM, Csf1 contributed to interferon-responsive activities, and regulated the cellular responses to interleukin-1 (IL-1) family members in mononuclear phagocytes (Fig. 5E and Supplementary Fig. 6A-C). These findings underscore the conserved regulatory mechanisms of FAPs on other cell types during regeneration.

### Resolving subpopulations of SCs and mononuclear phagocytes dependent on FAP signaling

#### FAPs and SC subpopulations

We reclustered just the SC parent population to obtain a high-resolution analysis of putative SC subpopulations. Five subpopulations with distinct features were identified (Fig. 6A-C), two of which appreciably expressed myogenic genes (Fig. 6C, clusters 1 and 2). Interestingly, cells in cluster 3, marked by immune-related gene expression and thus potentially an “immune SC” recently described^32^, were significantly diminished in depleted mice (Fig. 6A, B). Since myogenic genes were limited in expression, we also mapped these genes on the parent UMAP to interrogate spatial resolution in the SC cluster and found expression localized on the periphery (Supplementary Fig. 10A). We observed the SCs located towards the FAP and tenocyte clusters had minimal expression of myogenic genes (Supplementary Fig. 10A, B). This led us to further interrogate the identity of these cells at higher resolution, reclustering the SC, FAPs, and tenocyte grouping together. In this refined clustering, FAPs and tenocytes maintained close proximity in the dimensionally reduced space across their subclusters, whereas SCs were distributed among distinct clusters (Supplementary Fig. 10C, D). Two SC subclusters expressed myogenic genes and corresponded to the previously identified clusters 1 and 2. Additional subclusters, while identified as SCs based on total gene expression, also expressed marker subsets associated with pericytes, Schwann cells, immune cells, and an unidentified phenotype originating from the main SC cluster and tapering off towards the FAPs and tenocytes clusters of the original UMAP. While the reclustering of the parent grouping of 3 populations led to further resolution of the SC population, the initial clustering demonstrates the “intersection” cells remain more like SCs than other cell types in the regenerative milieu, and reveals a potential phenotypic variation in this population, one that will be further investigated in future studies.

**Figure 6.**
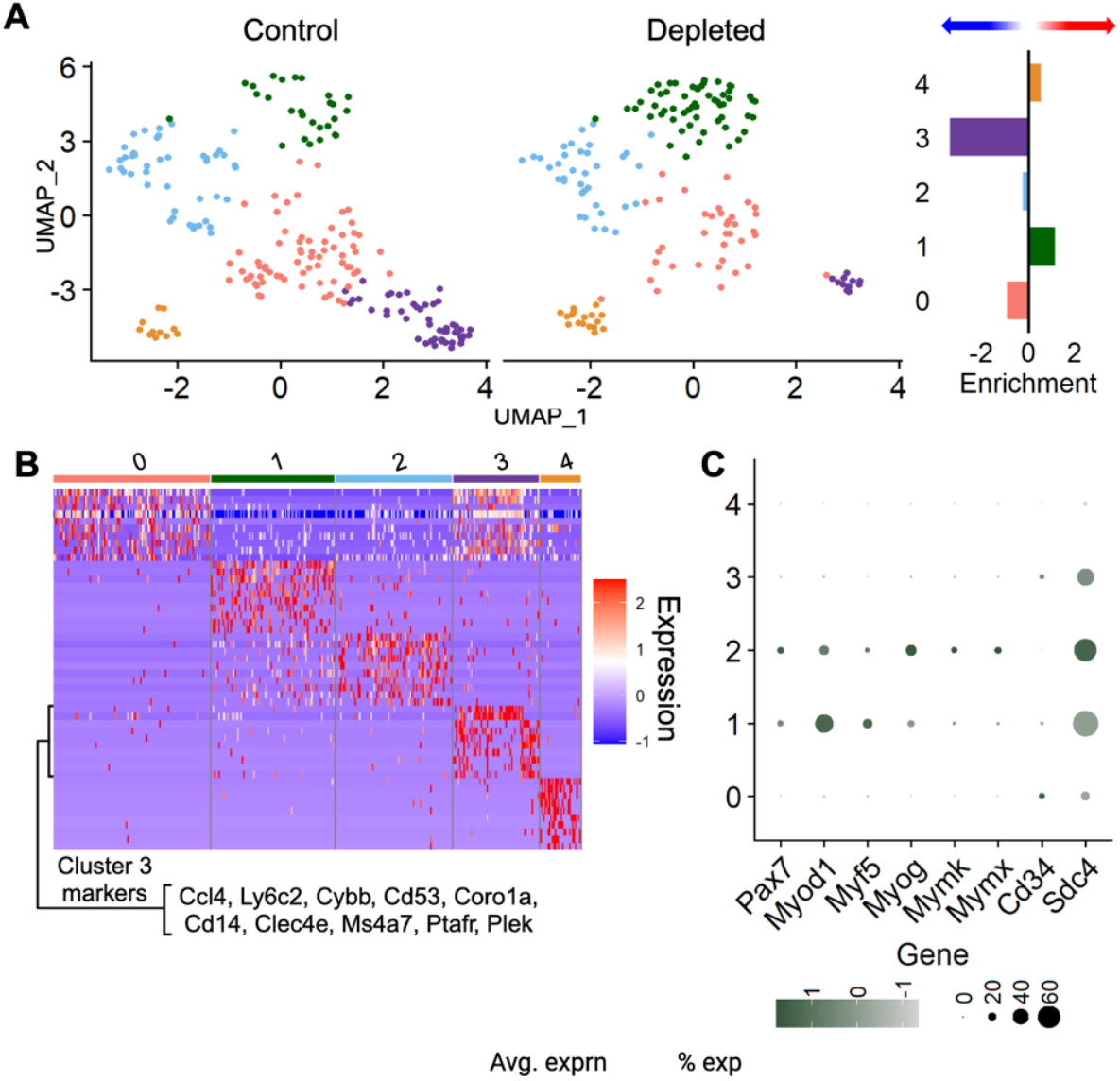
Molecular influence of FAPs on subpopulations of SCs in injured SkM. (A) UMAPs of SCs independently clustered from Fig. 4A; cells are colored by subcluster. Enrichment scores of each subcluster in SCs are graphed, with positive values indicating an enrichment in FAP-depleted mice and negative values indicating an enrichment in control mice. (B) Heatmap showing the top 10 enriched features in each subpopulation in SCs resolved with the FindAllMarkers function. Markers of cluster 3 are defined due to this population being the major cluster impacted by FAP-depletions (See graph in “D.” (C) A dot plot representing the abundance and expression level of myogenic genes in SC subpopulations.

#### FAPs and mononuclear phagocyte subpopulations

Given that the mononuclear phagocytes are known to be heterogeneous, performing complex tasks during muscle repair, this population was reclustered. We identified four distinct subsets (Fig. 7A). Two of these subsets displayed significant features corresponding to pro-inflammatory and anti-inflammatory macrophage phenotypes, respectively. At the same time, another subgroup exhibited high levels of MHC II molecules, indicating a role in antigen presentation (Supplementary Fig. 12 D, E). Interestingly, a Mφ subset was characterized by profound expression of Gpnmb (termed Gpnmb+ Mφ, Fig. 7A, and S12D, E), a population previously described as promoting muscle regeneration^33^. This subset shared features with the regenerative Mφ subset, but lacked expression of Mrc1, which encodes the anti-inflammatory macrophage surface marker CD206 (Supplementary Fig. 12D, E). CD206+ macrophages are stimulated by anti-inflammatory cytokines such as IL-10 and TGF-β and have been described as pro-regenerative in healing human and mouse SkM. However, a recent study reported that CD206+ macrophages can also play a role in hindering muscle regeneration indirectly by regulating the secretion of follistatin from FAPs^34–39^. Therefore, the context-dependent functions of CD206+ macrophages are yet to be thoroughly investigated.

**Figure 7.**
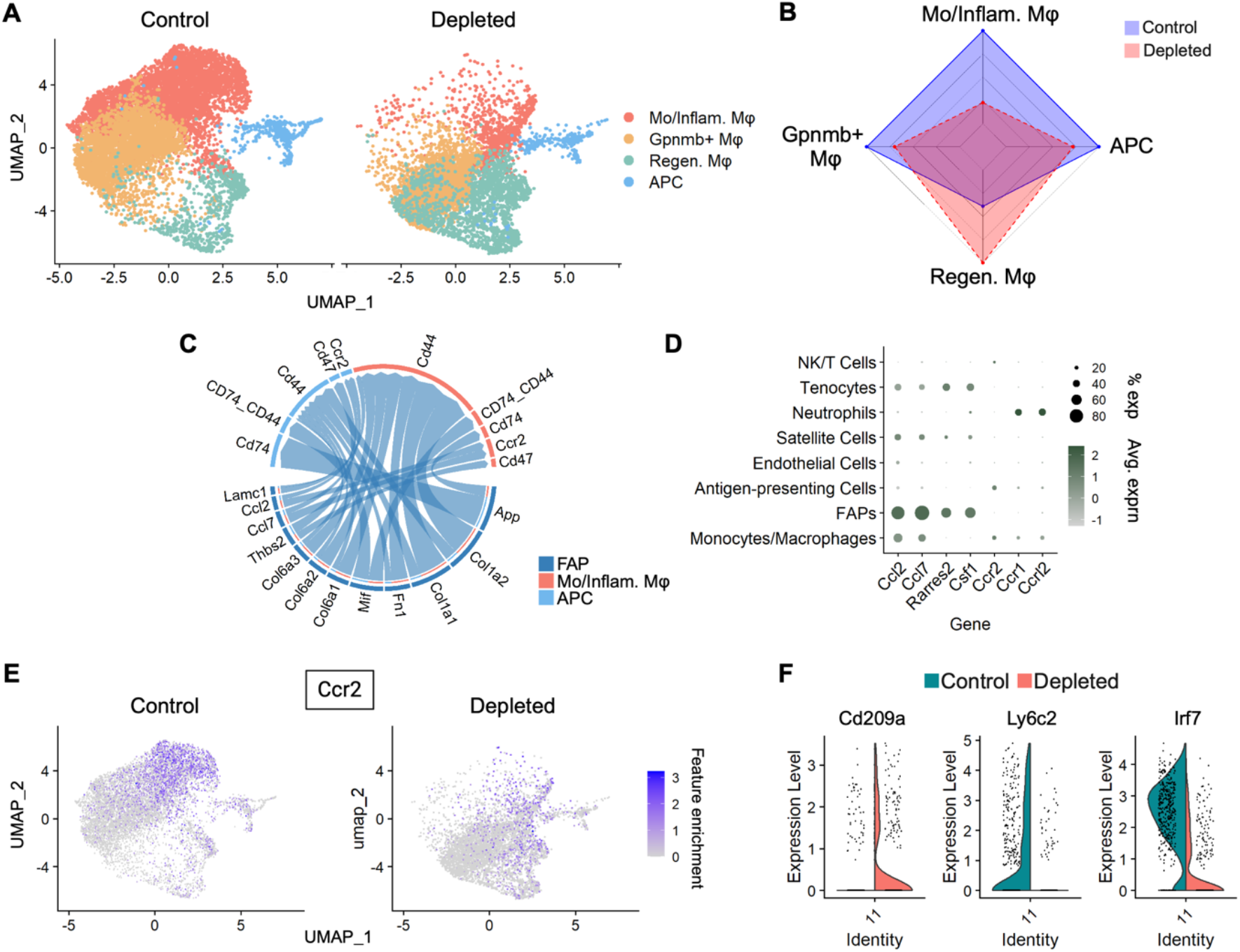
Molecular influence of FAPs on subpopulations of mononuclear phagocytes in injured SkM. (A) UMAPs of mononuclear phagocytes after independently reclustering from Fig. 4A. Cells are colored by noted subpopulation identity. (Inflam. - Inflammatory; Regen. - Regenerative) (B) Radar plot showing the difference in the number of each subpopulation in “C,” comparing FAP-depleted versus control samples. (C) Interactions between FAPs (sender) and Mo/APCs (receiver). The thickness of lines represents interaction strength. (D) Dot plot representing the abundance and expression level of genes associated with chemotaxis (except for Csf1) in mononuclear phagocytes across cell populations in SkM at 3 dpi. (E) UMAPs of mononuclear phagocytes overlaid with the expression level of Ccr2 in the noted samples. (F) Violin plots representing the expression levels of noted genes in APCs, split by sample.

In FAP-depleted mice, the inflammatory myeloid population decreased by 78%, while Gpnmb+ macrophages and APCs saw reductions of 30% and 28%, respectively. Interestingly, the regenerative macrophage subset expanded by 157% (Fig. 7B). Following injury, monocyte populations are typically recruited to injured tissue from circulation and either remain monocytes or differentiate into pro-inflammatory macrophages. We used CellChat to investigate the potential FAP-dependent chemotaxis mechanisms involved in this recruitment process. This analysis revealed that FAP expression of the chemoattractants Ccl2 and Ccl7, target Ccr2 on inflammatory mononuclear phagocytes and APCs, potentially explaining the reduced infiltration and irreversible decline in their numbers when FAPs are depleted (Fig. 7C-E). Additionally, Ccl7 can interact with Ccr1 on neutrophils (Fig. 7D and Supplementary Fig. 12B), so the loss of FAPs as its source seemingly results in a reduced number of neutrophils in FAP-depleted mice.

Interestingly, we noted that Rarres2 (encoding the ligand chemerin) is almost uniquely expressed in FAPs in the regenerative scenario (Fig. 5D). While our NicheNet analyses did not detect a strong ability of Rarres2 to regulate DEGs in mononuclear phagocytes (Fig. 5D), a subset of myeloid progenitors and neutrophils expressed its receptor, Ccrl2 (Supplementary Fig. 12F). Furthermore, Ccrl2-expressing monocytes, macrophages, and neutrophils were appreciably reduced in FAP-depleted mice, which indicates the potential for lack of corresponding ligands derived from FAPs being the cause (Supplementary Fig. 12F). CellChat analyses suggest that FAP-derived Csf1 can target Csf1r on Gpnmb+ macrophages, signaling that has been reported to be crucial for the self-renewal of tissue-resident macrophages^23^. These data indicate that the observed reduction in Gpnmb+ macrophages in FAP-depleted mice is directly related to signaling from FAPs (Fig. 7D and Supplementary Fig. 12G, H). Additionally, the Csf1-Csf1r signaling between FAPs and regenerative macrophages accentuates the pleiotropic signaling originating from FAPs to mediate the regenerative macrophage response. We also noted a subfraction of the regenerative macrophages expressing cell cycle-associated genes, indicating active proliferation (Supplementary Fig. 12I). Moreover, with FAPs depleted, a higher number of proliferative Mrc1+ macrophages were present when compared to control (Supplementary Fig. 12J), suggesting a compensatory increase in proliferation for the expansion of the macrophage population. Finally, we noted the APCs in depleted mice were mainly committed to a dendritic cell fate, while those in control mice retained their monocyte identity (Fig. 7F). These data further support the role of FAPs in regulating the immune cascade in SkM regeneration, as mature dendritic cell presence is dysregulated when FAPs are depleted, presenting earlier at 3 dpi then in control mice.

## Discussion

FAPs are essential for maintaining and regenerating SkM, primarily by modulating the environment to support local stem cells and immune cells. However, the specific regulatory factors secreted by FAPs and their full impact on intracellular signaling remain unclear. This gap in knowledge stems from the challenge of deciphering complex, multicellular signaling networks in both homeostatic and regenerative contexts.

In this study, we used scRNA-seq to capture a comprehensive view of mononuclear cells in homeostatic and regenerative SkM. We integrated this with advanced bioinformatics to infer FAP-dependent signals that influence other cells, shaping their roles in SkM maintenance and regeneration. However, these predictions are correlative and require systematic validation, a reductionist approach that limits the assembly of broad signaling networks. To overcome this, we combined scRNA-seq inference with FAP-depleted datasets from both uninjured and injured SkM, enabling us to assess the impact of FAPs on predicted mechanisms at a systems level. This approach mapped the FAP interactome across cellular states, uncovering a multifaceted web of intercellular signaling. Our dataset serves as a resource to predict FAP-dependent communication and test whether inferred mechanisms are functionally linked to FAPs at a multicellular level, an achievement not yet realized. Additionally, we identified new FAP-mediated regulatory mechanisms in homeostatic and regenerative SkM, while also confirming several recently reported pathways.

### FAPs in Homeostasis

Our scRNA-seq-based cellular profiling in homeostasis, using control and FAP-depleted input, revealed the impact of FAPs on cell number and composition with greater resolution than FACS profiling. We observed a decrease in Mo/Mφs and B cells, along with a slight increase in neutrophils, NK/T cells, and Schwann cells. SC numbers remained unchanged, consistent with previous findings showing only long-term effects of FAP depletion in homeostasis^12^. Despite stable SC numbers, pathway analysis indicated that adhesion and location maintenance processes were downregulated upon FAP depletion, underscoring the role of FAPs in supporting SCs and stabilizing their niche. Further DEG analyses identified significant FAP-dependent gene expression changes in SCs, particularly in Tgfb2, multiple collagens, MMP2, and Timp2. Interaction analyses linked these effects to the surface expression of Sdc4 and Cd44, genes known to regulate SC activation, proliferation, migration, and differentiation through ECM interactions^40–43^. SC reclustering further revealed the impact of FAPs on subcluster representation, reinforcing their role in maintaining the SC niche. While additional studies are needed to fully dissect this mechanism, myogenic gene expression and gene ontology analyses implicate connective tissue (ECM) as a key regulator.

Pathway-resolved signaling (PRS, Supplementary Fig. 4) reveals ECM-receptor interactions as a major aspect of FAP-mediated signaling in homeostasis. As expected, SCs are a primary target, but PRS analyses also include monocytes and macrophages (Mo/Mφs), which show significant DEG changes following FAP depletion. Notably, Sdc4 and Cd44 are identified as key regulators of FAP-Mo/Mφ interactions. Further analyses reveal that FAP-derived Csf1 and Vegfa significantly impact Mo/Mφs, supporting previous findings and validating our dataset^23,44^. CSF1 from FAPs has been shown to sustain the self-renewal of tissue-resident macrophages in homeostatic muscle^23^. We demonstrate that Csf1 and Vegfa, predominantly expressed in FAPs, may regulate Mo/Mφ activity by modulating miRNA transcription and processing. While miRNA expression is broadly implicated in SkM regeneration and disease, its role in macrophages within uninjured muscle remains unclear. After acute injury, BM-derived monocyte miR-223-3p helps balance inflammatory responses in SkM^45^, but the relevance of miRNA regulation in homeostasis warrants further study. Following Mo/Mφ reclustering, we observed that the absence of ligands regulating miRNA transcription correlates with a reduced number of IFN-responsive Mo/Mφs, though their functional significance remains unexplored. These findings suggest that FAPs help sustain Mo/Mφ responsiveness to endogenous and exogenous stimuli. While circulating Mo/Mφ contributions cannot be entirely excluded, our data reveal an intriguing role for FAPs in maintaining Mo/Mφ representation in homeostatic muscle.

We identified an interaction between FAPs and neutrophils in homeostatic muscle. Neutrophils have a short half-life, circulating in the bloodstream for only a few hours before undergoing apoptosis^46^. While mature and immature neutrophils are present in peripheral blood, they are typically not tissue-resident under normal conditions. However, we observed a cluster of neutrophils in uninjured muscle. Previous studies have shown that non-perfused mice retain detectable granulocytes in muscle, whereas perfused mice do not, suggesting a circulatory origin^19,47^. We analyzed FAP-neutrophil interactions and found evidence suggesting that FAPs contribute to neutrophil maturation. This interaction is predicted to be mediated through Cd44 and Cxcr2, implicating ECM involvement and chemotaxis, respectively. Furthermore, we identified a secondary effect of FAP depletion through reciprocal signaling between neutrophils and Mo/Mφs, mediated by adhesion ligands and chemokines. These findings highlight the complex multicellular effects of FAPs, both by directly influencing cell populations and indirectly shaping cellular responses through primary interactions.

### FAPs in Regeneration

Our results effectively resolved the impact of FAPs on the cellular composition of the regenerative milieu at 3 dpi, when the regenerative mononuclear cell response peaks in SkM. FAP depletion led to reduced numbers of SCs, Mo/Mφs, NK cells, neutrophils, and APCs, while tenocytes slightly increased. While most of these changes confirm previous FACS-based profiling, our approach uniquely dissected immune cell populations beyond the pan-marker CD45, and revealed a FAP-dependent effect on putative tenocytes. Additionally, we observed a decrease in IFN-β signaling across multiple cell populations in FAP-depleted SkM, suggesting that FAPs are required for global immune signaling and essential for effective regeneration. Though the role of IFN-β in muscle regeneration remains unexplored, our findings implicate FAPs as a potential source. Notably, this broad, cell-type-specific impact would have been undetectable without the FAP-depleted dataset.

Our injured SkM dataset captured sufficient SC numbers for scRNA-seq analyses, overcoming a common limitation in 10X scRNA-seq at 3 dpi, where SCs are often underrepresented, presumably due to technical challenges in processing. With adequate SC representation, our results revealed a close relationship between SCs, tenocytes, and FAPs. Reclustering these three populations showed that tenocytes and FAPs remained distinct, while SCs fractionated into multiple subpopulations. Curiously, two myogenic marker-expressing subclusters contained ∼30% more cells in the depleted dataset than in controls, whereas the remaining SC subclusters were more prevalent in controls. This suggests that FAPs are required for SC subcluster identity, and in their absence, SC genetic diversity is restricted.

Further analysis of FAP-SC interactions identified Fgf7, Il6, Il33, and Fbn1 as FAP-dependent regulators of SC function. While collagens are not exclusive to FAPs in the regenerative milieu, collagens and Fn1 are predicted to interact with Sdc4 and Cd44 on SCs. This suggests that FAP-derived collagens and Fn1 may be necessary to maintain sufficient levels in the regenerative environment. Recent findings in a porcine SkM screen and extrapolated to mouse and human SkM support FGF7 as a FAP-expressed ligand that targets myogenic cells to promote proliferation^48^. Similarly, Il6 signaling, partially originating from FAPs, has been described as a potent inducer of myogenesis early in the regenerative milieu^22^, though persistently high IL-6 levels are linked to chronic inflammation, muscle weakness, and atrophy^49–51^. Interestingly, we also identified FAP-derived Fgf7 and Il6 signaling affecting tenocytes and endothelial cells, emphasizing the pleiotropic role of FAPs as master regulators of the regenerative milieu. Collectively, these findings validate the robustness of our dataset, reinforce recent reports of FAP-dependent signaling, and reveal an extensive FAP-driven network influencing SCs, tenocytes, and endothelial cells during regeneration.

We also reclustered SCs alone to assess their genetic diversity and subcluster heterogeneity. Similar to the SC-tenocyte-FAP reclustering, this analysis identified five SC subpopulations, with two retaining the majority of myogenic gene expression. Among them, cluster “3” showed the greatest change, exhibiting a reduction in numbers upon FAP depletion. Hierarchical gene expression analysis revealed that this cluster expressed several immune-related markers ^52–56^, suggesting similarity to the recently identified “immune SC” population. While the function of these cells remains unclear, further investigation could reveal an interesting role for these cells in SkM regeneration.

FAP depletion had a profound impact on Mo/Mφ populations, as evident in control and depleted UMAPs. Our approach provided enhanced resolution by profiling the entire Mo/Mφ population, uncovering a complex signaling network between FAPs and Mo/Mφs. An initial observation was the reduction of Mo/Mφs expressing markers associated with Mo and inflammatory Mφs, likely mediated by FAP-derived Ccl2/7 and Rarres2 and their corresponding receptors (Ccr2 and Ccrl2, respectively) on Mo/Mφs. Ccl2/7-Ccr2 chemotaxis is well-established across tissues, with a recent finding linking senescent FAPs in aged homeostatic tissue to monocyte recruitment and phenotypic transition, which are associated with harmful collagen deposition and fibrosis^24^. Our study implicates these pathways in regeneration following acute injury in young animals. Following FAP depletion, we also observed a decrease in Ccr1 and Ccrl2-expressing neutrophils, reinforcing the far-reaching impact of FAP interactions. While a reduction in inflammatory cells was expected, our data also reveal dysregulation in Mφs with a regenerative phenotype. Surprisingly, regenerative Mφs increased, but we also observed a decrease in Gpnmb+ Mφs, which share markers with regenerative Mφs and are associated with the pro-regenerative phenotype. Whether Gpnmb+ Mφs are a distinct subset or functionally related to regenerative Mφs remains unclear. FAPs are predicted to regulate both regenerative macrophage populations through collagens, Fn1, and Csf1, with their respective receptors (Cd44, Sdc4, and Csf1r) expressed on Gpnmb+ and regenerative Mφs. Additionally, FAPs express Il-33 within 12 hours post-injury^57^, a chemokine that promotes the transition to pro-regenerative Mφs essential for SkM repair. By 3 dpi, our data show that tenocytes are the primary source of Il-33, while FAP expression is minimal, highlighting the temporal modulation of Il-33 in the multicellular regenerative milieu.

In conclusion, we have constructed a FAP interactome model by validating predicted interactions in control conditions and testing their functional relevance through FAP depletion. This approach reinforces FAPs as master regulators of both homeostatic and regenerative SkM environments, orchestrating a multicellular signaling network. Although FAP depletion was systemic, many identified pathways align with recently published findings, reinforcing the robustness of our dataset and methodology. Furthermore, our work shows that many FAP-derived signals are shared between homeostasis and regeneration, while also uncovering new pathways and cellular responses that specifically depend on FAP signaling. By leveraging a systems-level approach, we mapped intercellular networks that reductionist methods alone could not resolve, providing the field with a comprehensive FAP interactome that advances our understanding of SkM regulation.

## Acknowledgments

We would like to thank current and past members of the Wosczyna laboratory for experimental input and discussion. We would also like to thank the members of the Stanford University School of Medicine Genome Sequencing Service Center for Chromium 10X processing and single-cell RNA sequencing.

Our work was supported by the NIA/NIH (AG053438 to MNW) and the Grossman School of Medicine to MNW.

## Contributions

XL, EEPC, YCH, CSM, and MNW designed the experiments and analyses. XL, EEPC, YCH, BWG, CSM, and MNW performed the experiments and analyses. XL, EEPC, SAB, and MNW wrote the manuscript.

## Methods

### Animal models

Mice in this study were bred on a C57BL/6J (Jackson Labs 000664) background. All mice were male, and ages ranged from 3-6 months. Mice were housed in rooms on a 12-hour on/off cycle, with food and water provided ad libitum. The genotypes of experimental mice referred to as depleted were PDGFRα^CreER/+^; R26^DTA/NG^ or R26^DTA/+ 12^ and controls were littermates that did not carry both the CreER and DTA. All mice received tamoxifen to control for toxicity. Tamoxifen (Sigma-Aldrich) was resuspended in corn oil (Sigma-Aldrich) at a concentration of ∼0.1 mg/gm and administered via intraperitoneal (IP) injection. Mice were dosed 7 times over 9 days, with recovery days on days 4 and 7. Following dosing, mice were rehoused in tamoxifen-free cages and subjected to a 9-day chase period prior to injury. The mice were maintained according to the Institutional Animal Care and Use Committee (IACUC)-approved protocols of the Veterans Affairs Palo Alto Health Care System (VAPAHCS) and the New York University Grossman School of Medicine.

### BaCl_2_ Injury

Intramuscular injury of the lower hindlimb muscles was performed on mice anesthetized with isoflurane. Multiple injections of 1.2% BaCl_2_ solution (Sigma-Aldrich) using an insulin needle were split across lower hindlimb muscles, including the tibialis anterior (TA) and gastrocnemius muscles. Injections were followed with multiple punctures throughout the lower hindlimb using the same needle to help distribute the BaCl_2_ and achieve consistent injuries. Contralateral hindlimbs were used as uninjured controls. Ethiqa XR was given by subcutaneous injection (0.05-0.1 mg/kg) to control pain.

### Cell isolation

SkM was harvested from either the uninjured or injured lower hindlimb. Harvested muscles were minced mechanically with scissors and then digested in a 760 U/mL Collagenase Type II enzyme (Worthington Biochemical) solution at 37°C in a shaking water bath for one hour. Wash media (Ham’s F10 (Cytiva) + 10% horse serum (ThermoFisher Scientific)) was used to neutralize the digest before resuspending the samples in a second enzyme solution containing Collagenase Type II (stock 1000U/ml, and Dispase (stock 11U/ml, ThermoFisher Scientific), both at a 1:8.5 dilution of stock, at 37°C in a shaking water bath for 30 minutes. Samples were syringed through a 21G needle before being diluted with wash media and then passed through a 40um filter to remove larger debris. Cell viability was assessed with propidium iodide (PI), and negative cells were submitted to the Stanford University School of Medicine Genomics core for 10X Chromium (version 2) processing and sequencing on an Illumina HiSeq 4000 sequencer. Two biological replicates were sequenced per experimental condition. For cellular profiling by FACS, following the 40 um filter step, cells were stained with Pacific Blue-Sca-1 (BioLegend 108120), PE/Cyanine7-CD106 (BioLegend 105720), APC-CD31 (BioLegend 102510), APC-CD45 (BioLegend 103112), or PE/Cyanine7-CD31 (Invitrogen #25-0311-82), all at 1:100 dilution for 30 minutes on a shaker at 4^°^C. Gating strategies are shown in Supplementary Fig. 2C and D. Samples were processed using a Sony MA900 Multiapplication Cell Sorter for cell profiling and a BD Biosciences FACS Aria III for cell sorting for sequencing.

### Single-cell RNAseq processing workflow

Cell Ranger (v 6.0.2) was used to obtain the raw expression counts for samples by mapping them to the reference transcriptome GRCm3. In short, the binary base call sequence file was converted to FASTQ format using the Cell Ranger mkfastq function; the count matrices and quality control metrics were generated with the Cell Ranger count function, and results from multiple samples were merged using the Cell Ranger aggr function. Seurat R package (v 5.0.3)^25^ was used to process the Cell Ranger output in RStudio (v 4.3.2). Quality control was conducted to exclude low-quality cells, specifically those with mitochondrial content exceeding 20% or extreme feature counts (fewer than 200, indicating potential debris, or more than 2,500, suggesting possible doublets). Only genes detected in more than three cells were retained. Feature expression was normalized using default settings, and all features were scaled. The top 2,500 features showing high cell-to-cell variation were identified for downstream analysis. Principal component analysis (PCA) was then performed to reduce data dimensionality, computing 30 PCs, with the first 10 to 17 used for future clustering analysis. Finally, cells were clustered using the k-nearest neighbors (k-NN) algorithm at a resolution of 0.4 to 1.0. The result of dimensional reduction was visualized using UMAP, a non-linear dimensionality reduction algorithm with the ‘euclidean’ metric selected in the RunUMAP function to measure the spatial distance between data points. Clusters with similar marker expressions were consolidated into a single group. The number variation of each cell population between FAP-depleted and control datasets was visualized using the ggplot function in ggplot2 R package (v 3.5.1)^58^ and the radarchart function in fmsb R package (v 0.7.6). The FindAllMarkers function was used to identify the most enriched features that are expressed in more than 25% of each cluster. The FindMarkers function was used to determine the differentially expressed genes (DEGs) between conditions, with the default Wilcox test. The Benjamini & Hochberg method was used to calculate the adjusted p-value using the p.adjust function in the stats R package (v 4.3.2). Only genes with log_2_ fold change (log_2_FC) ≥ 0.25 and adjusted p-value ≤ 0.05 were taken into account. DEGs were plotted using the EnhancedVolcano function in the EnhancedVolcano R package (v 1.20.0). Populations of interest were reclustered, and visualization/analysis was performed using the same methods described above.

### Neutrophil trajectory analysis

Cells identified as neutrophils in the homeostasis dataset were extracted and transformed from a Seurat object to a large cell_data_set object using the embedded as.cell_data_set function in monocle3 (v 1.0.0)^28^. The dataset was pre-processed using default settings. The first 50 principal components were calculated, and the dataset was subjected to dimensional reduction and visualization through UMAP. Cells were clustered using Leiden algorithm-based community detection. A principal graph was learned to construct the trajectory that the cells take through spatially. Cells previously characterized as “proliferating neutrophils” were manually assigned as the root and ordered along the trajectory. Throughout all Monocle 3 plots, the cells retained their original labels from the Seurat processing.

### Cell-cell interaction interference - CellChat

Seurat objects containing cells from the control datasets were transformed into large CellChat objects using the embedded createCellChat function in the CellChat R package (v 2.1.1)^27^. Individual ligand-receptor interaction databases categorized by mode (ECM-receptor, secreted signaling, cell-cell interactions) or a combination of modes were used for cell-cell communication prediction. The analysis and visualization were operated with default parameters.

### Ligand-target prediction - NicheNet

The nichenetr R package (version 2.0.4)^26^ was used to study the regulatory mechanisms of FAP-derived signals in other cell populations. In a single analysis, only one receiver population from either homeostasis or regeneration datasets was investigated. In brief, features expressed in over 10% of FAPs are considered ligand candidates, and those expressed in over 10% of receiver cells are considered receptor candidates. Only ligand candidates that pair with receptor candidates through the ligand-receptor network are retained for further analysis. Gene in receiver cells differentially expressed between FAP-depleted and control (adjusted p-value < 0.05, calculated by the Benjamini & Hochberg method; abs(log_2_FC) > 0.25) and expressed in at least 5% of receiver cells in either control or FAP-depleted were used as gene sets of interest. A ligand-target matrix showing the potential regulatory strength of ligands for target genes was used for ligand activity prediction. The top 10 prioritized ligands and their predicted target genes that are in the gene sets of interest (DEGs) and among the top 120 known targets for each prioritized ligand, were visualized by a heatmap using the make_heatmap_ggplot function in nichenetr. The expression of the ligands across all detected cell populations in muscles was visualized by the DotPlot function in Seurat.

### Biological pathway enrichment analyses - clusterprofiler and GSEA

The enrichGO function in the clusterProfiler R package (v 4.10.1)^59^ was used for enrichment analysis on gene sets. A gene set generated from one of the following was assessed in a single analysis: a) DEGs between FAP-depleted and control in a cell population (abs(log_2_FC) > 0.5; expressed in at least 10% of cells in this population in either control or FAP-depleted), identified using the FindMarkers function and divided into upregulated (log_2_FC > 0) and down-regulated (log_2_FC < 0) groups for independent assessments; b) target genes in the receiver cell population predicted to be regulated by a specific ligand from FAPs (in NicheNet analysis); c) the top50 features most enriched in each subpopulation after reclustering. Analyses were carried out using the “BP” subontology in the genome-wide annotation org.Mm.eg.db, with the keytype of input gene specified as “SYMBOL”. The top 10 enriched biological pathways in each analysis were visualized by the ggplot function.

Gene set enrichment analysis (GSEA) was used as an alternative approach for functional enrichment assessment. The analysis was conducted using GSEA software (v 4.3.3)^60,61^ on the FAP-depleted and control datasets in regeneration condition, with the comparison of each gene reported as signal-to-noise ratio and used for ranking. The M5 ontology gene sets in MSigDB were used for enrichment analysis, focusing on the Biological Process subset. The analysis used the gene set permutation without collapse to gene symbols. GO terms whose FDR < 0.05 were considered significantly enriched.

### Graph generation and statistical analysis

GraphPad Prism software (v10) was used to generate graphs and assess statistics. The arithmetic mean of each experimental group is displayed in graphs, with error bars illustrating the standard error of the mean (SEM). The number of replicates (n) used in each experiment is specified in figure legends. Unpaired t-tests were performed to calculate statistical significance, with a significance threshold set at a p-value of ≤ 0.05. Statistics for sequencing analyses are noted in the above corresponding sections. FACS data were analyzed and visualized using FlowJo software (v 10.9.0).

## Ethics declarations

The authors declare no competing interests.

## Abbreviations

SkM: skeletal muscle
MSC: mesenchymal stem cell
PACE: PDGFRα^CreER^
DTA: diphtheria toxin A
FACS: fluorescence-activated cell sorting
scRNA-seq: single-cell RNA-sequencing
DEG: differentially expressed gene
CT: control
DP: FAP-depleted
FAP: fibroadipogenic progenitor
MuSC/SC: muscle stem cell/satellite cell
EC: endothelial cell
IC: immune cell
Teno: tenocyte
Neut: neutrophil
NK cell: natural killer cell
Sch: Schwann cell
SMC: smooth muscle cell
Mo: monocyte
Mφ: macrophage
APC: antigen-presenting cell
Unk: unknown
ECM: extracellular matrix
UMAP: uniform manifold approximation and projection
PRS: pathway-resolved signaling

