## Supplemental Figures 1-13 and Tables 1-3 for "Mesenchymal Stem Cell-Mediated Intercellular Communication: Mapping the Interactome for Skeletal Muscle Homeostasis and Regeneration"

### Supplementary figures

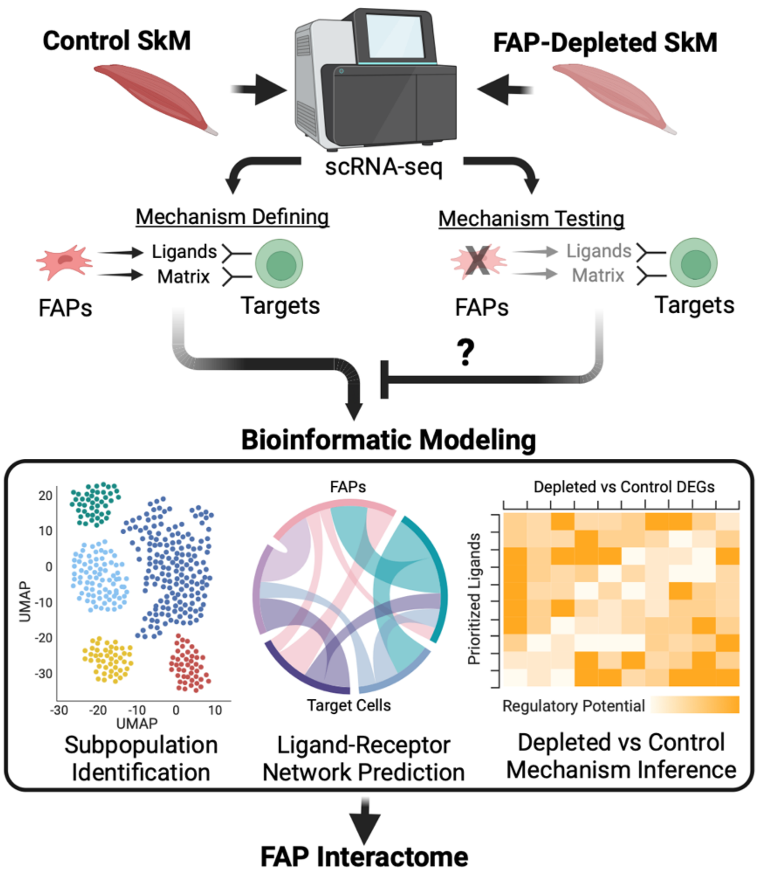


**Supplementary Figure 1. Graphical Abstract.**

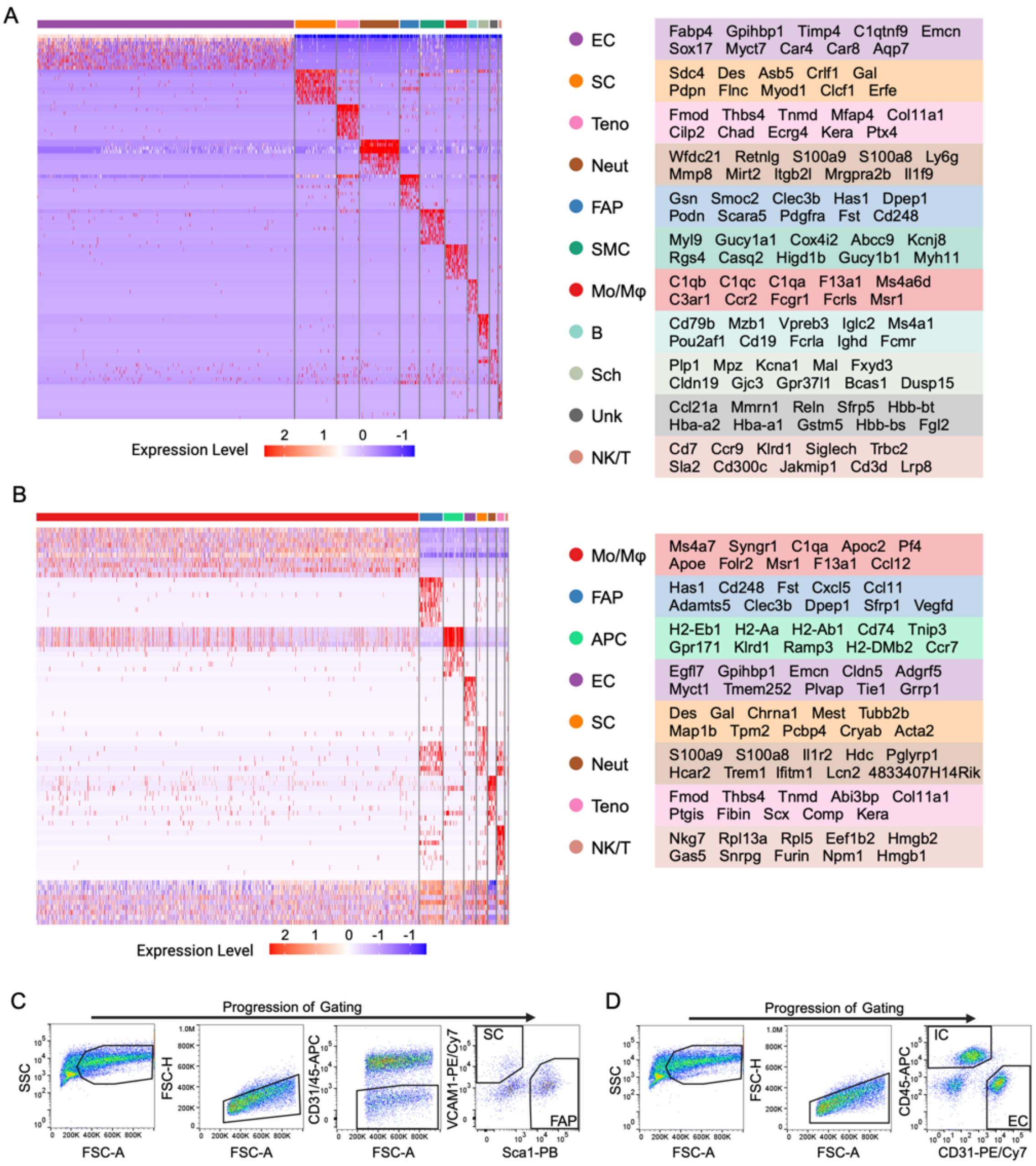


**Supplementary Figure 2. Identifying cellular populations in homeostatic and regenerating SkM.** (A, B) Heatmaps showing the top 10 enriched features in each population of (A) uninjured and (B) injured SkM resolved with FindAllMarkers function. Markers are noted by the corresponding color to the populations in the heatmap. ECs - endothelial cells, SC - satellite cells, Teno - tenocytes, Neut - neutrophils, FAPs - fibroadipogenic progenitors, SMC - smooth muscle cells, Mo/Mφs - monocytes/macrophages, B - B cells, Sch - Schwann cells, NK/T - NK cells/T cells, APC - antigen-presenting cells, Unk - unknown. (C, D) Representative gating strategy for identifying cellular fractions in SkM. IC - immune cell.

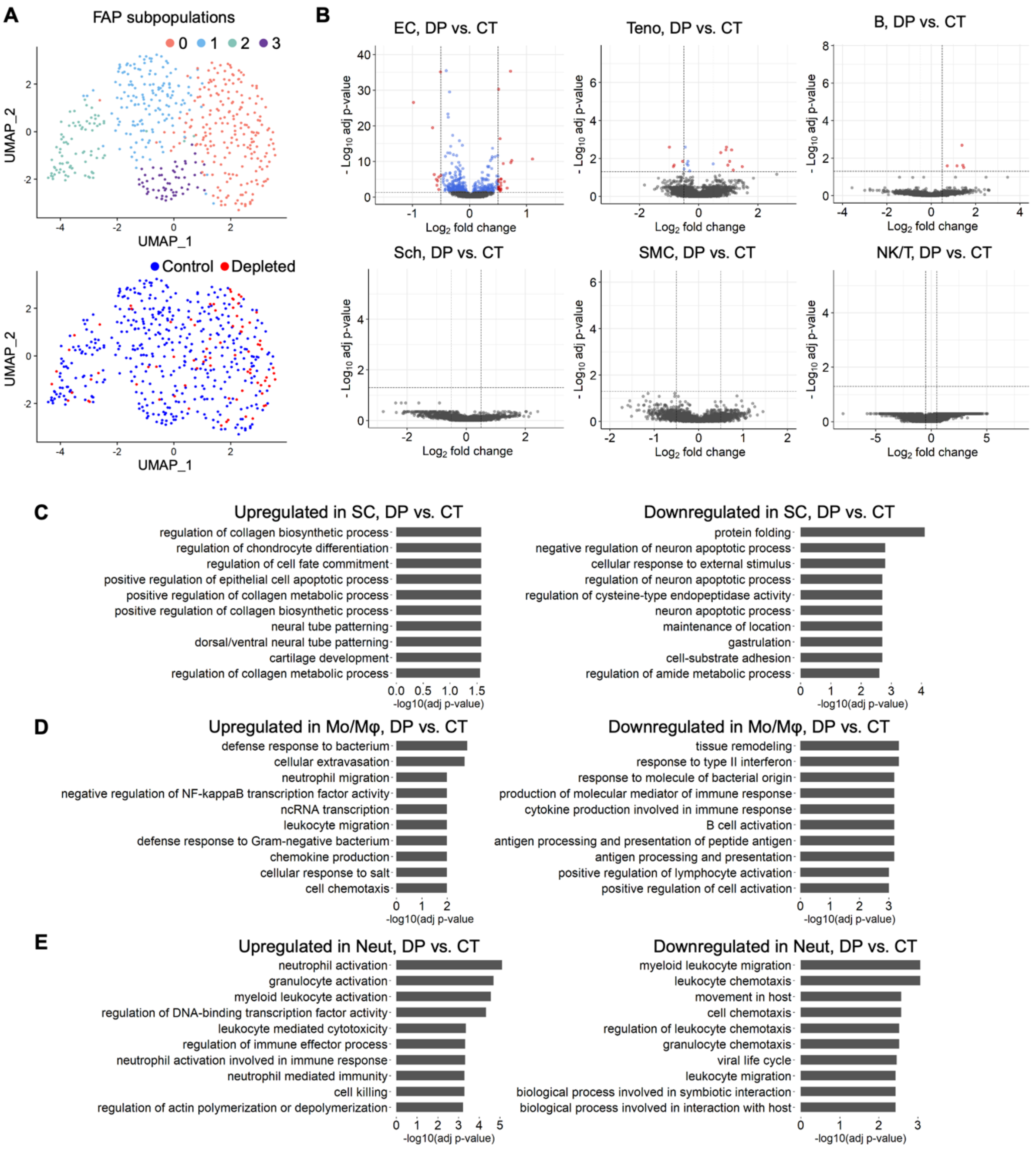


**Supplementary Figure 3. Elucidating FAP-dependent gene expression and molecular pathways in cells of the homeostatic niche.** (A) UMAPs of FAPs after independently reclustering from Fig. 2A. Cells are colored by noted subpopulation identity (top) or sample (bottom). (B) Volcano plots representing DEGs in the noted cell populations from control and FAP-depleted samples. Red: DEGs with $log_2FC > 0.5$ and adjusted p-value < 0.05. Blue: DEGs with $log_2FC \leq 0.5$ and adjusted p-value < 0.05. Gray: DEGs with adjusted p-value ≥ 0.05. (C-E) Top 10 upregulated and downregulated biological processes in noted cell populations in uninjured SkM, comparing FAP-depleted versus control samples. DP - FAP-depleted, CT -

control; FAPs - fibroadipogenic progenitors, EC - endothelial cells, Teno - tenocytes, B - B cells, Sch - Schwann cells, SMC - smooth muscle cells, NK/T - natural killer/T-cells, SCs - satellite cells, Mo/Mφs - monocytes/macrophages, Neut - neutrophils.

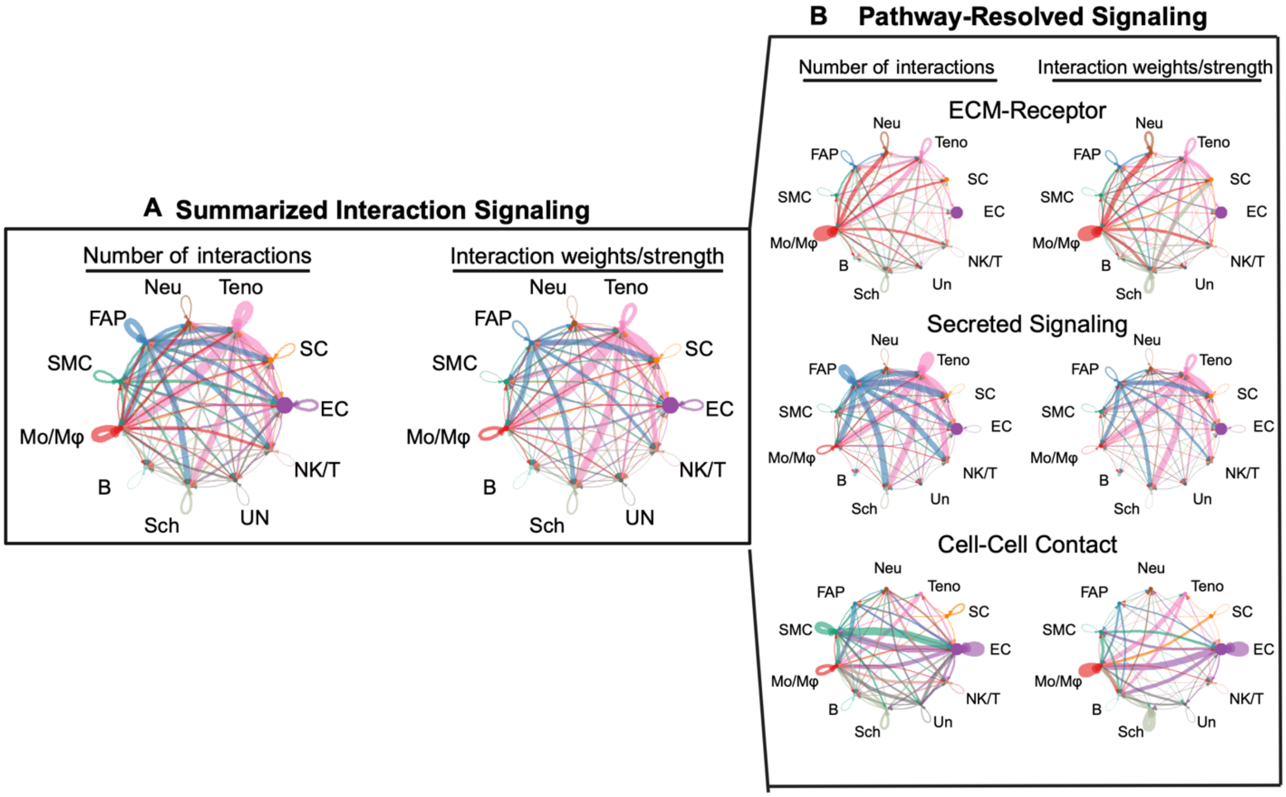


**Supplementary Figure 4. FAP interactions in homeostatic SkM**. (A, B) Chord plots from CellChat analyses summarize the interactions between cells of homeostatic SkM. (A), and resolved by pathway types (B). The thickness of the lines represents the number of interactions (left) and interaction weights/strength (right).

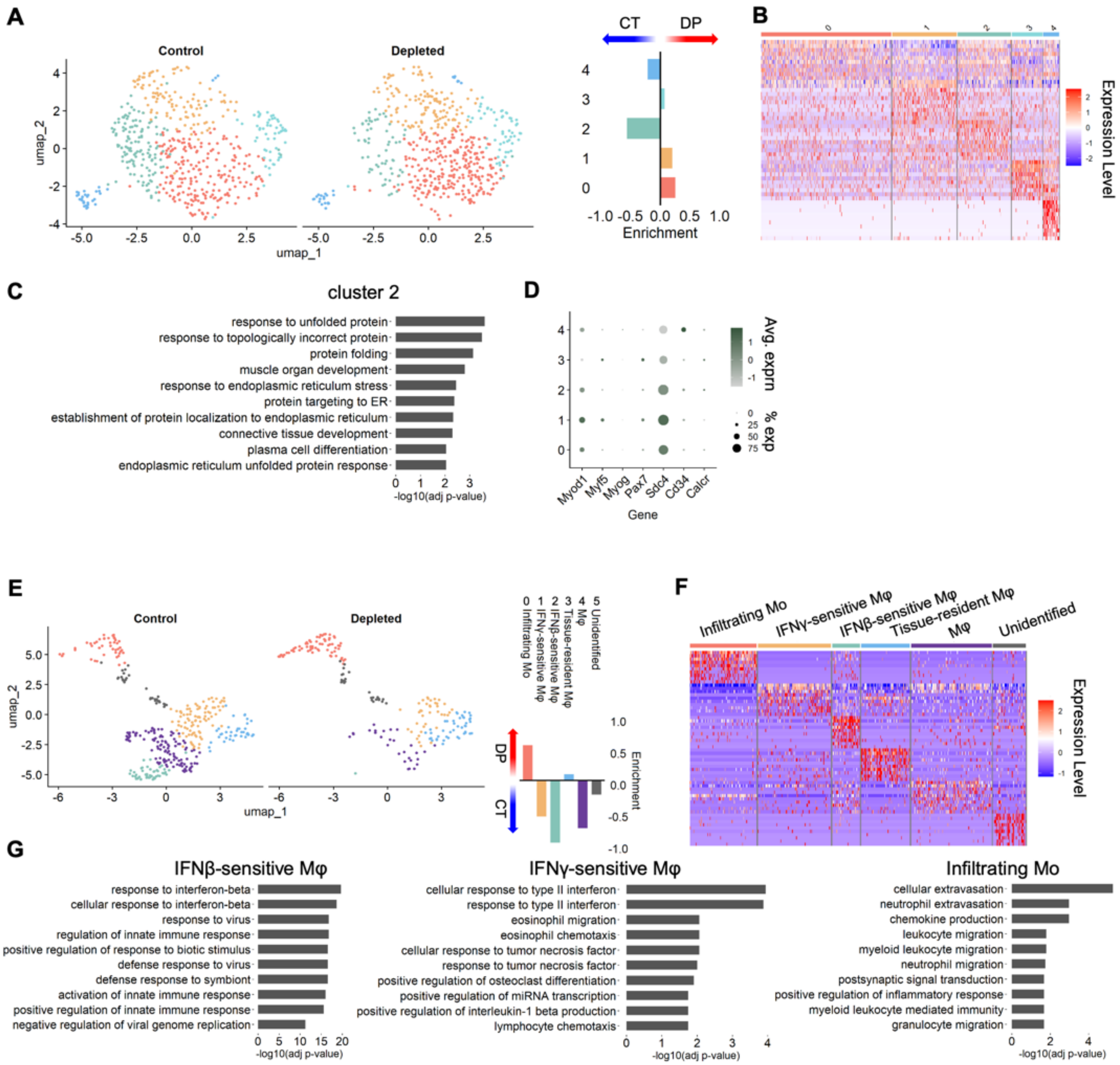


**Supplementary Figure 5. Molecular influence of FAPs on subpopulations of SCs and Mo/Mφs in SkM homeostasis.** (A) UMAPs of SCs after independent reclustering of the parent population in Fig. 2A. Subpopulations are color-coded, corresponding with colors in the enrichment score plot (right) of SC subclusters in uninjured SkM. Positive scores represent a greater abundance of cells in FAP-depleted mice, while negative scores indicate a greater abundance in control mice. (B) Heatmap showing the top 10 enriched features in each (A) subpopulation in SCs in uninjured SkM resolved with the FindAllMarkers function. Specific markers are listed in Supplementary Table 1. (C) Top 10 biological processes in SC cluster 2, compared to other SC subclusters. (D) Dot plot representing the expression of genes associated with myogenesis across SC populations in uninjured SkM. (E) UMAPs of Mo/Mφs after independently reclustering their parent population from Fig. 2A. Subpopulations are color-coded, corresponding to their noted identity in the enrichment score plot of Mo/Mφ subclusters in uninjured SkM (right). Positive scores represent a greater abundance of cells in FAP-depleted mice, while negative scores indicate a greater abundance in control mice. (F) Heatmap showing the top 10 enriched features in each subpopulation of Mo/Mφs in uninjured SkM resolved with the FindAllMarkers function. Specific markers are listed in Supplementary Table 2. (G) Top 10 biological enriched processes in noted Mo/Mφ subpopulations versus other subpopulations.

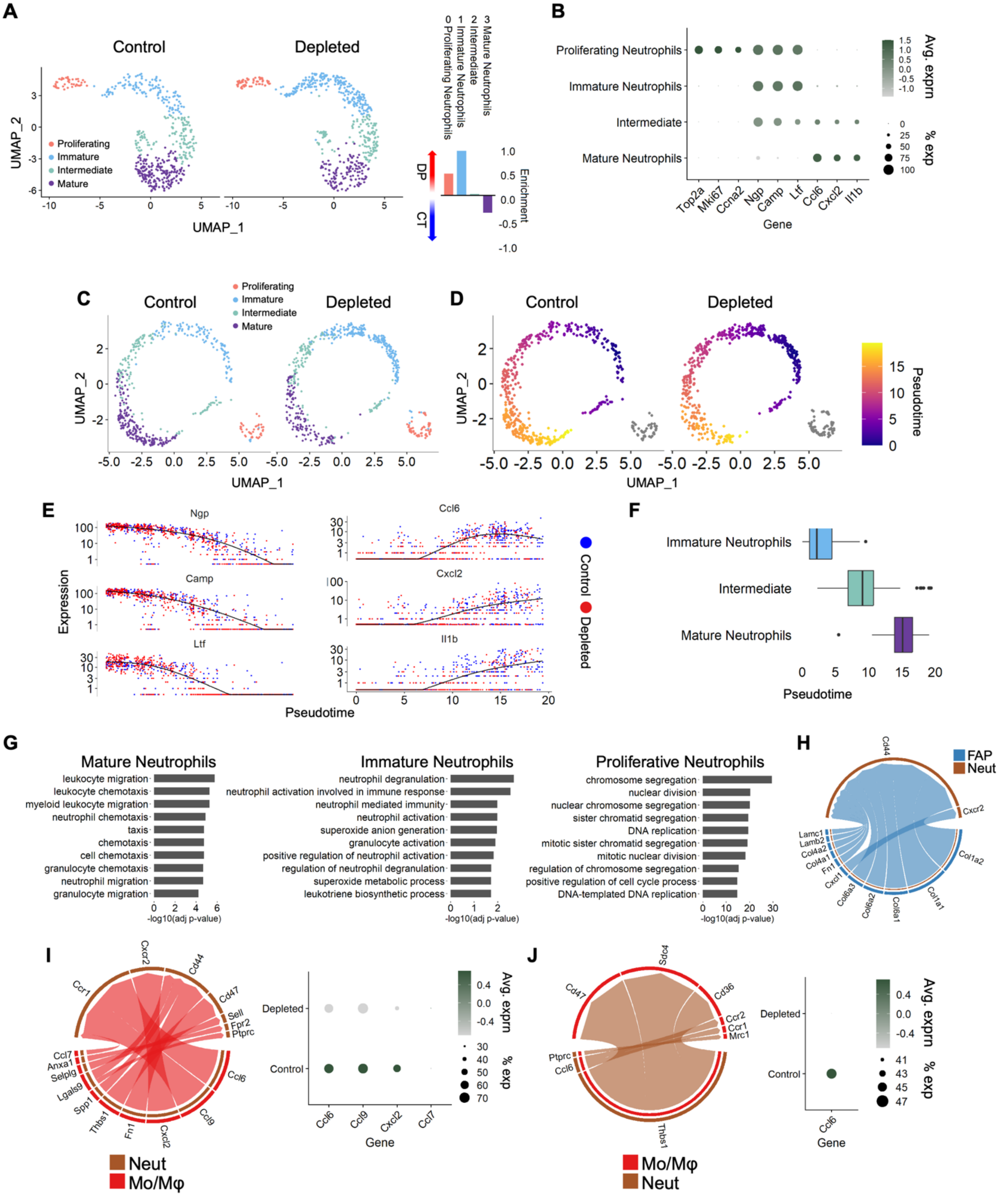

**Supplementary Figure 6. Molecular impact of FAPs on neutrophil subpopulations in homeostasis.** (A) UMAPs of neutrophils after independently reclustering from their parent population in Fig. 2A. Cells are color-coded by subpopulation identity, with corresponding the enrichment score plot of neutrophils subclusters in uninjured SkM (right). Positive scores represent a greater abundance of cells in FAP-depleted mice, while negative scores indicate a greater abundance in control mice. (B) Dot plot representing the abundance and expression level of genes associated with neutrophil developmental stages across neutrophil populations in uninjured SkM. (C) UMAPs of neutrophils are created by Monocle 3, with cells colored by the neutrophil developmental stage. (D) UMAPs of neutrophils created by Monocle 3, with cells colored by pseudotime. Immature neutrophils were manually set as the root of the main trajectory. Most proliferating neutrophils were in a minor separated trajectory (gray). (E) Expression of marker genes indicating neutrophil developmental stage along pseudotime. (F) Abundance of neutrophil subpopulations along pseudotime. (G) Top 10 biological processes in noted neutrophil subpopulations, compared to others. (H) Interactions between FAPs (sender) and neutrophils (receiver). The thickness of lines represents interaction strength. (I) Left: Interactions between Mo/Mφs (sender) and neutrophils (receiver). The thickness of lines represents interaction strength. Right: Dot plot representing the abundance and expression level of genes associated with chemokines in Mo/Mφs across samples in uninjured SkM. (J) Left: Interactions between neutrophils (sender) and Mo/Mφs (receiver). The thickness of lines represents interaction strength. Right: Dot plot representing the abundance and expression level of genes associated with chemokines in neutrophils across samples in uninjured SkM.

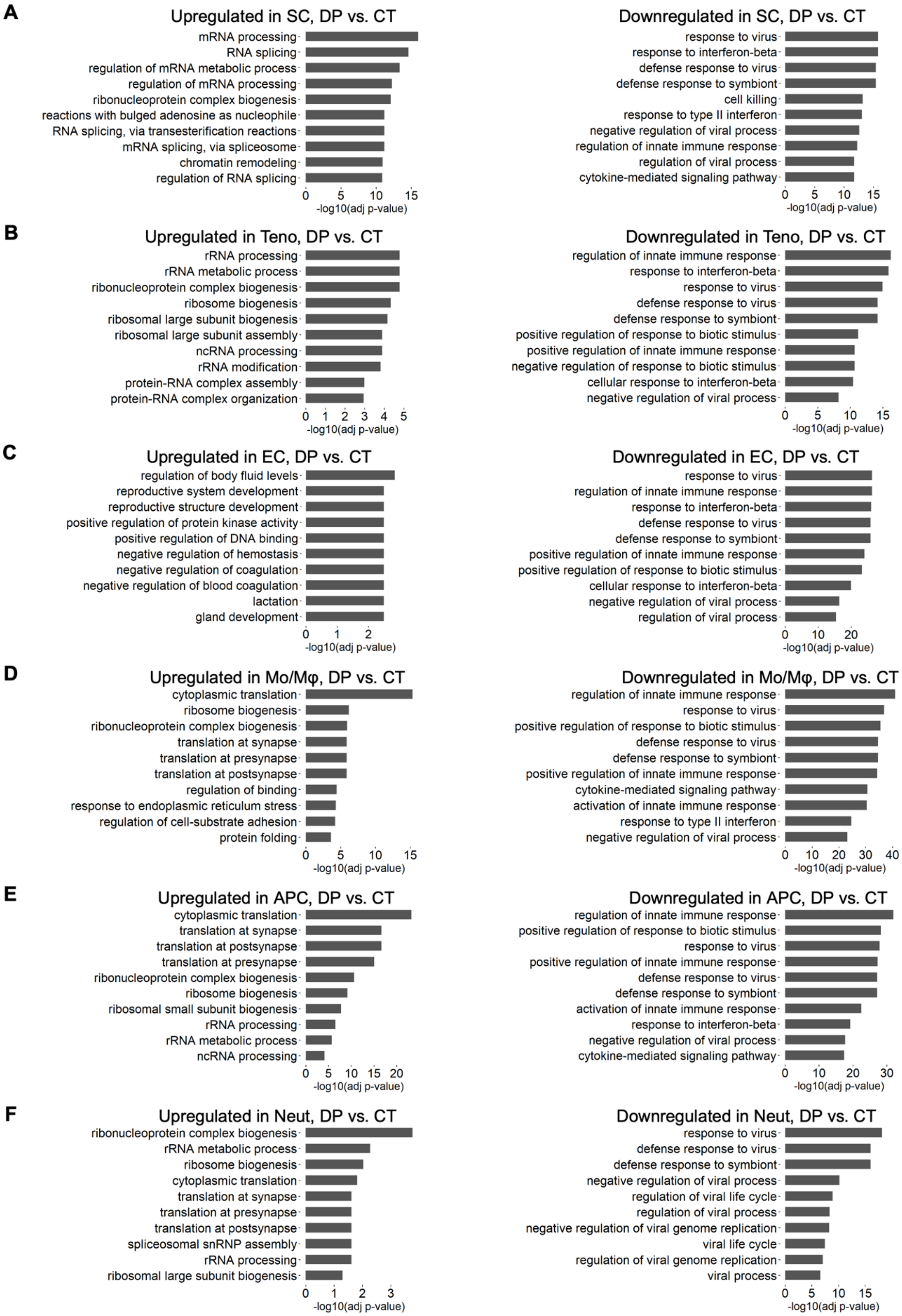
A
Upregulated in SC, DP vs. CT
mRNA processing
RNA splicing
regulation of mRNA metabolic process
regulation of mRNA processing
ribonucleoprotein complex biogenesis
reactions with bulged adenosine as nucleophile
RNA splicing, via transesterification reactions
mRNA splicing, via spliceosome
chromatin remodeling
regulation of RNA splicing
0 5 10 15
-log10(adj p-value)
Downregulated in SC, DP vs. CT
response to virus
response to interferon-beta
defense response to virus
defense response to symbiont
cell killing
response to type II interferon
negative regulation of viral process
regulation of innate immune response
regulation of viral process
cytokine-mediated signaling pathway
0 5 10 15
-log10(adj p-value)
B
Upregulated in Teno, DP vs. CT
rRNA processing
rRNA metabolic process
ribonucleoprotein complex biogenesis
ribosome biogenesis
ribosomal large subunit biogenesis
ribosomal large subunit assembly
ncRNA processing
rRNA modification
protein-RNA complex assembly
protein-RNA complex organization
0 1 2 3 4 5
-log10(adj p-value)
Downregulated in Teno, DP vs. CT
regulation of innate immune response
response to interferon-beta
response to virus
defense response to virus
defense response to symbiont
positive regulation of response to biotic stimulus
positive regulation of innate immune response
negative regulation of response to biotic stimulus
cellular response to interferon-beta
negative regulation of viral process
0 5 10 15
-log10(adj p-value)
C
Upregulated in EC, DP vs. CT
regulation of body fluid levels
reproductive system development
reproductive structure development
positive regulation of protein kinase activity
positive regulation of DNA binding
negative regulation of hemostasis
negative regulation of coagulation
negative regulation of blood coagulation
lactation
gland development
0 1 2
-log10(adj p-value)
Downregulated in EC, DP vs. CT
response to virus
regulation of innate immune response
response to interferon-beta
defense response to virus
defense response to symbiont
positive regulation of innate immune response
positive regulation of response to biotic stimulus
cellular response to interferon-beta
negative regulation of viral process
regulation of viral process
0 10 20
-log10(adj p-value)
D
Upregulated in Mo/Mφ, DP vs. CT
cytoplasmic translation
ribosome biogenesis
ribonucleoprotein complex biogenesis
translation at synapse
translation at presynapse
translation at postsynapse
regulation of binding
response to endoplasmic reticulum stress
regulation of cell-substrate adhesion
protein folding
0 5 10 15
-log10(adj p-value)
Downregulated in Mo/Mφ, DP vs. CT
regulation of innate immune response
response to virus
positive regulation of response to biotic stimulus
defense response to virus
defense response to symbiont
positive regulation of innate immune response
cytokine-mediated signaling pathway
activation of innate immune response
response to type II interferon
negative regulation of viral process
0 10 20 30 40
-log10(adj p-value)
E
Upregulated in APC, DP vs. CT
cytoplasmic translation
translation at synapse
translation at postsynapse
translation at presynapse
ribonucleoprotein complex biogenesis
ribosome biogenesis
ribosomal small subunit biogenesis
rRNA processing
rRNA metabolic process
ncRNA processing
0 5 10 15 20
-log10(adj p-value)
Downregulated in APC, DP vs. CT
regulation of innate immune response
positive regulation of response to biotic stimulus
response to virus
positive regulation of innate immune response
defense response to virus
defense response to symbiont
activation of innate immune response
response to interferon-beta
negative regulation of viral process
cytokine-mediated signaling pathway
0 10 20 30
-log10(adj p-value)
F
Upregulated in Neut, DP vs. CT
ribonucleoprotein complex biogenesis
rRNA metabolic process
ribosome biogenesis
cytoplasmic translation
translation at synapse
translation at presynapse
translation at postsynapse
spliceosomal snRNP assembly
rRNA processing
ribosomal large subunit biogenesis
0 1 2 3
-log10(adj p-value)
Downregulated in Neut, DP vs. CT
response to virus
defense response to virus
defense response to symbiont
negative regulation of viral process
regulation of viral life cycle
regulation of viral process
negative regulation of viral genome replication
viral life cycle
regulation of viral genome replication
viral process
0 5 10 15
-log10(adj p-value)

**Supplementary Figure 7. FAP-dependent molecular pathways across cell populations in SkM regeneration.** (A-F) Top 10 upregulated and downregulated biological processes in noted cell populations in 3 dpi SkM, comparing FAP-depleted (DP) versus control (CT) samples. Truncated term in (A) “reactions with bulged adenosine as nucleophile” = RNA splicing, via transesterification reactions with bulged adenosine as nucleophile. DP - FAP-depleted, CT - control; SCs - satellite cells, Teno - tenocytes, EC - endothelial cells, Mo/Mφs - monocytes/macrophages, APC - antigen-presenting cells, Neut - neutrophils.

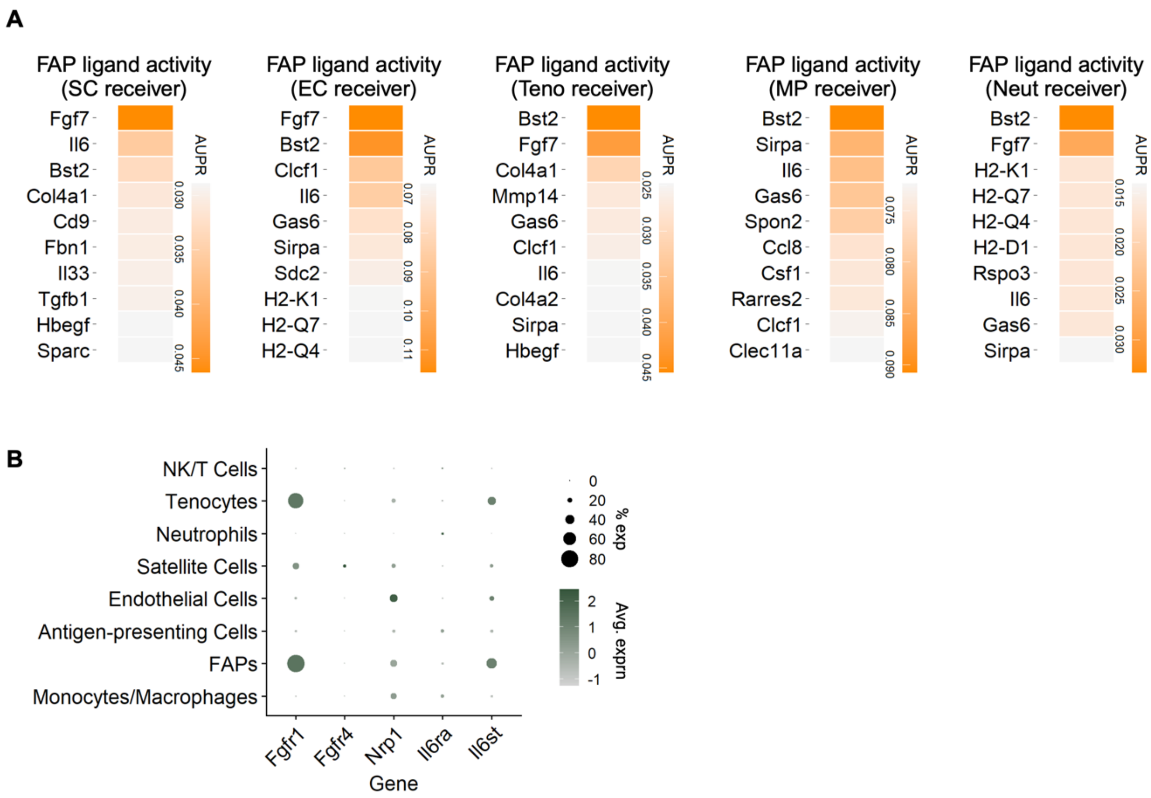


**Supplementary Figure 8. Regulatory potential of FAP-derived ligands for different cell populations in SkM regeneration.** (A) Top 10 FAP-derived ligand activity scores in 3 dpi SkM quantitated by the area under the precision-recall curve (AUPR), with noted cell populations set as receivers using NicheNet. (B) Dot plot representing the abundance and expression level of genes encoding receptors of FGF7 (Fgfr1, Fgfr4, Nrp1) and IL-6 (Il6ra, Il6st) across cell populations in SkM at 3 dpi.

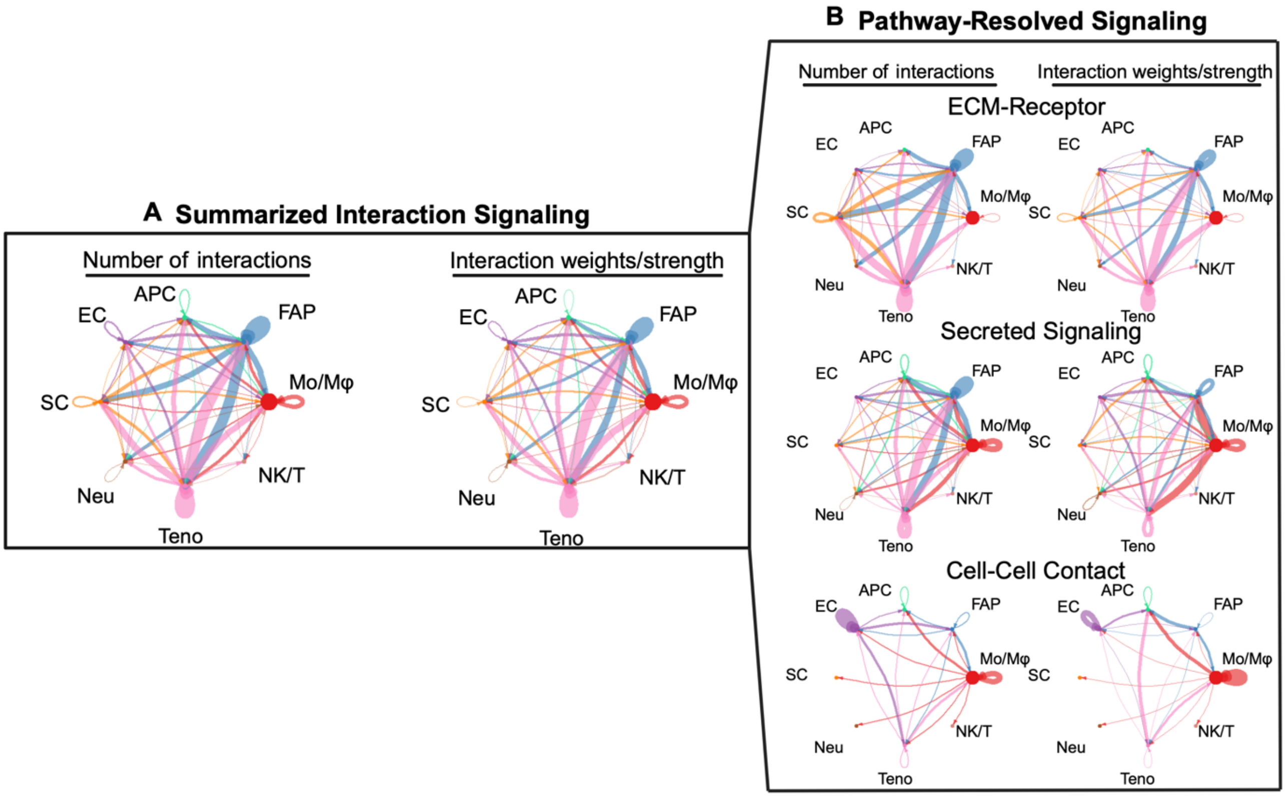


**Supplementary Figure 9. FAP interactions in regenerating SkM**. (A, B) Chord plots from CellChat analyses summarize the interactome between cells of regenerating SkM (A), and resolved by pathway types (B). The thickness of the lines represents the number of interactions (left) and interaction weights/strength (right).

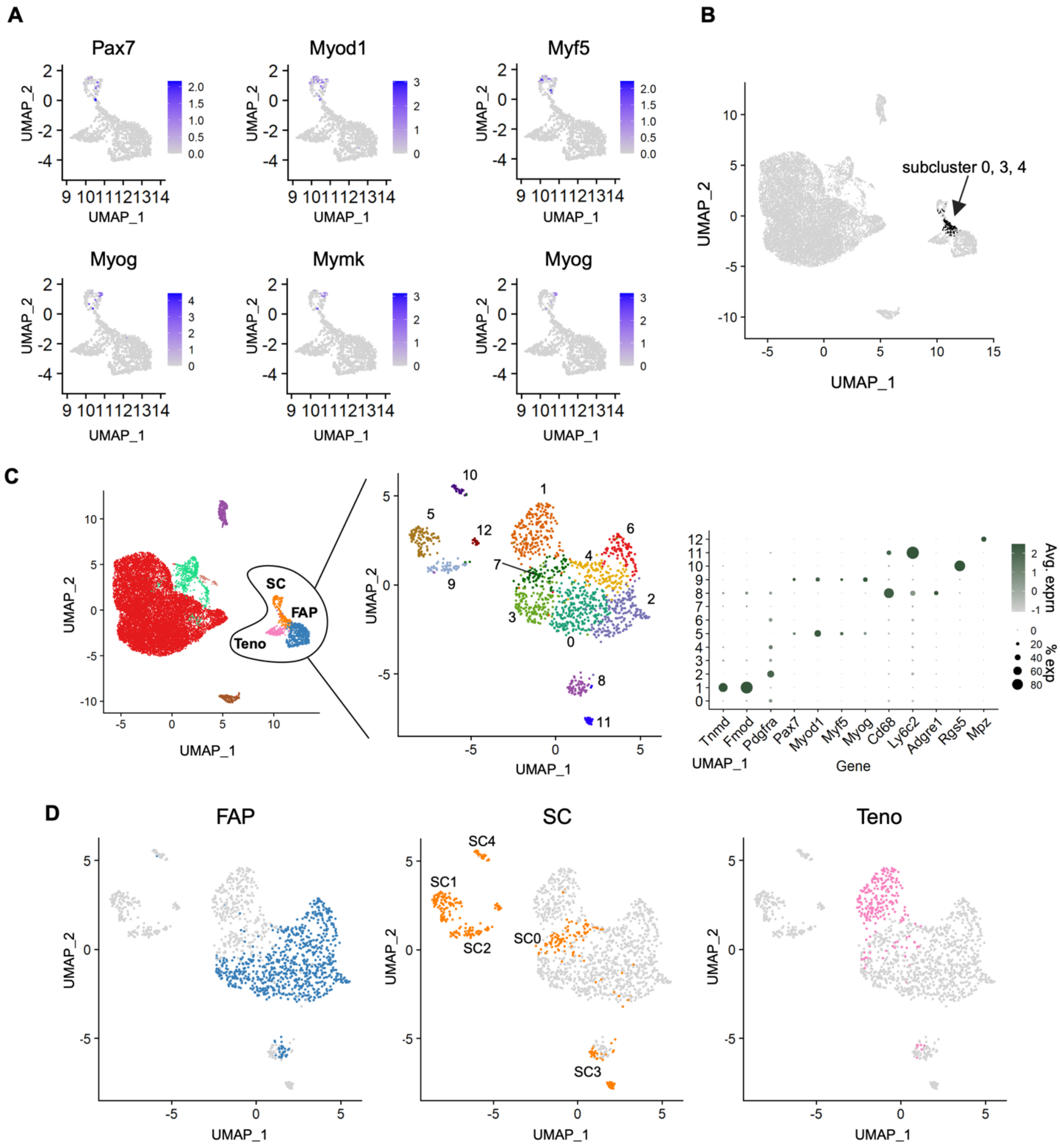


**Supplementary Figure 10. SC heterogeneity in SkM regeneration.** (A) UMAPs focused on the FAP, SC, and tenocyte populations in SkM regeneration (Fig. 4A) with the expression level of noted myogenic genes. Note, that FAP-depleted and control cells are overlaid in this representation. Purple gradient - feature enrichment score. (B) UMAP showing the distribution of cells in SC subclusters 0, 3, and 4 mapping to the parent UMAP from Fig. 4A. (C) UMAPs derived from scRNA-seq of cells from FAP-containing and depleted SkM at 3 dpi (left). UMAP of FAPs, SCs, and tenocytes after independently reclustering from the left panel, with cells colored by subcluster (middle). The dot plot (right) represents gene abundance and expression level for cluster identification across cell types in SkM at 3 dpi. (D) UMAPs of FAPs, SCs, and tenocytes after independently reclustering, with cells colored by their original identities in the parent UMAP (C, middle). SC subsets are labeled by features enriched and termed “like”.

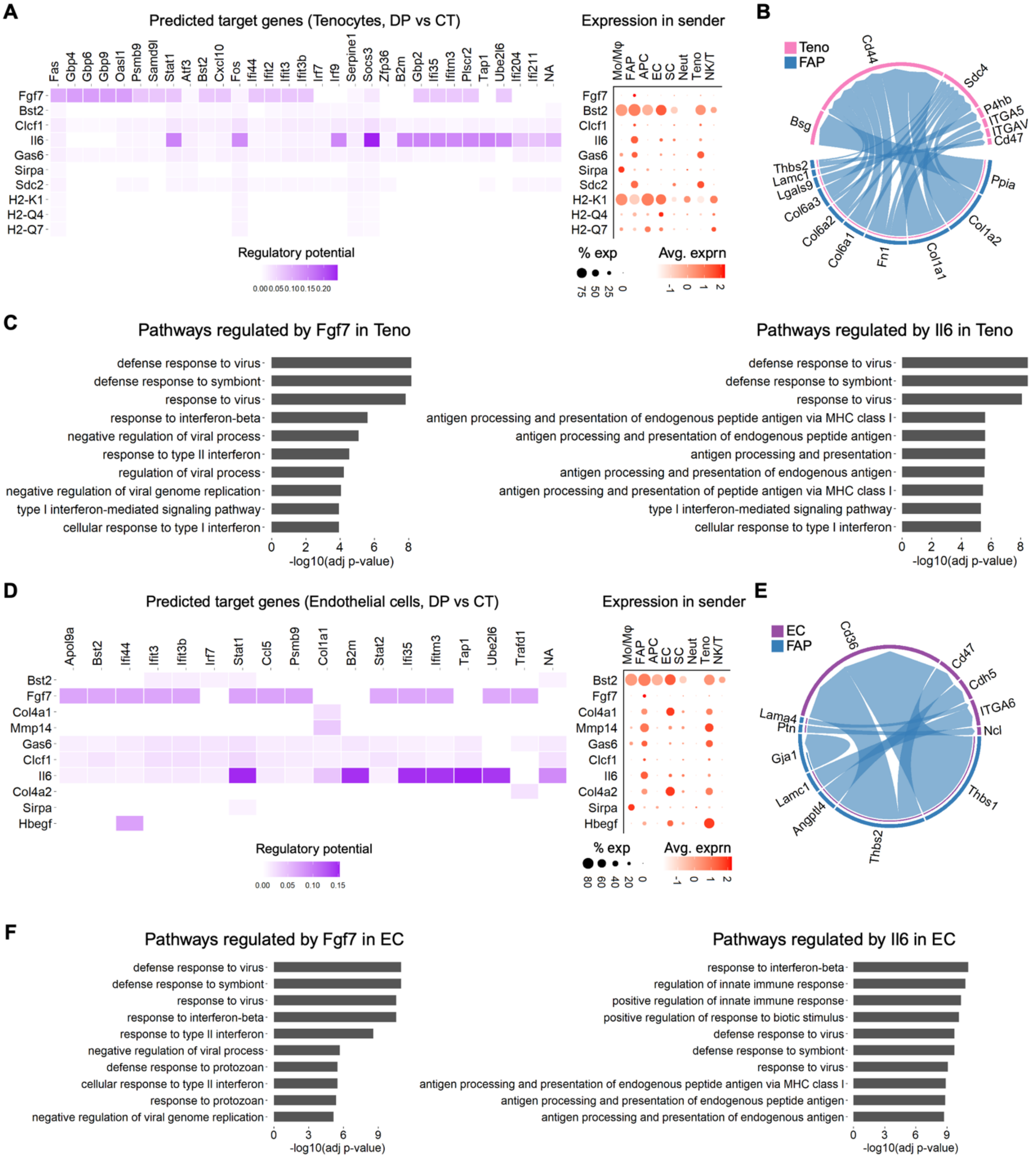

**Supplementary Figure 11. FAP-dependent regulatory mechanisms of tenocyte and endothelial cell activity in SkM regeneration.** (A) Top FAP-derived ligands with the highest activity and their predicted impact on DEGs in tenocytes, with the dot plot showing the abundance and expression level of these ligands across cell types in SkM at 3 dpi. (B) Interactions between FAPs (sender) and tenocytes (receiver). The thickness of lines represents interaction strength. (C) Top 10 biological processes in tenocytes inferred to be regulated by Fgf7 (left) and Il6 (right). (D) Top FAP-derived ligands with the highest activity and their predicted impact on DEGs in ECs, with the dot plot showing the abundance and expression level of these ligands across cell types in SkM at 3 dpi. (E) Interactions between FAPs (sender) and ECs (receiver). The thickness of lines represents interaction strength. (F) Top 10 biological processes in ECs inferred to be regulated by Fgf7 (left) and Il6 (right). Truncated terms in (B) and (E) "ITGA5" = ITGA5_ITGB1, "ITGAV" = ITGAV_ITGB1, and "ITGA6" = ITGA6_ITBG1.

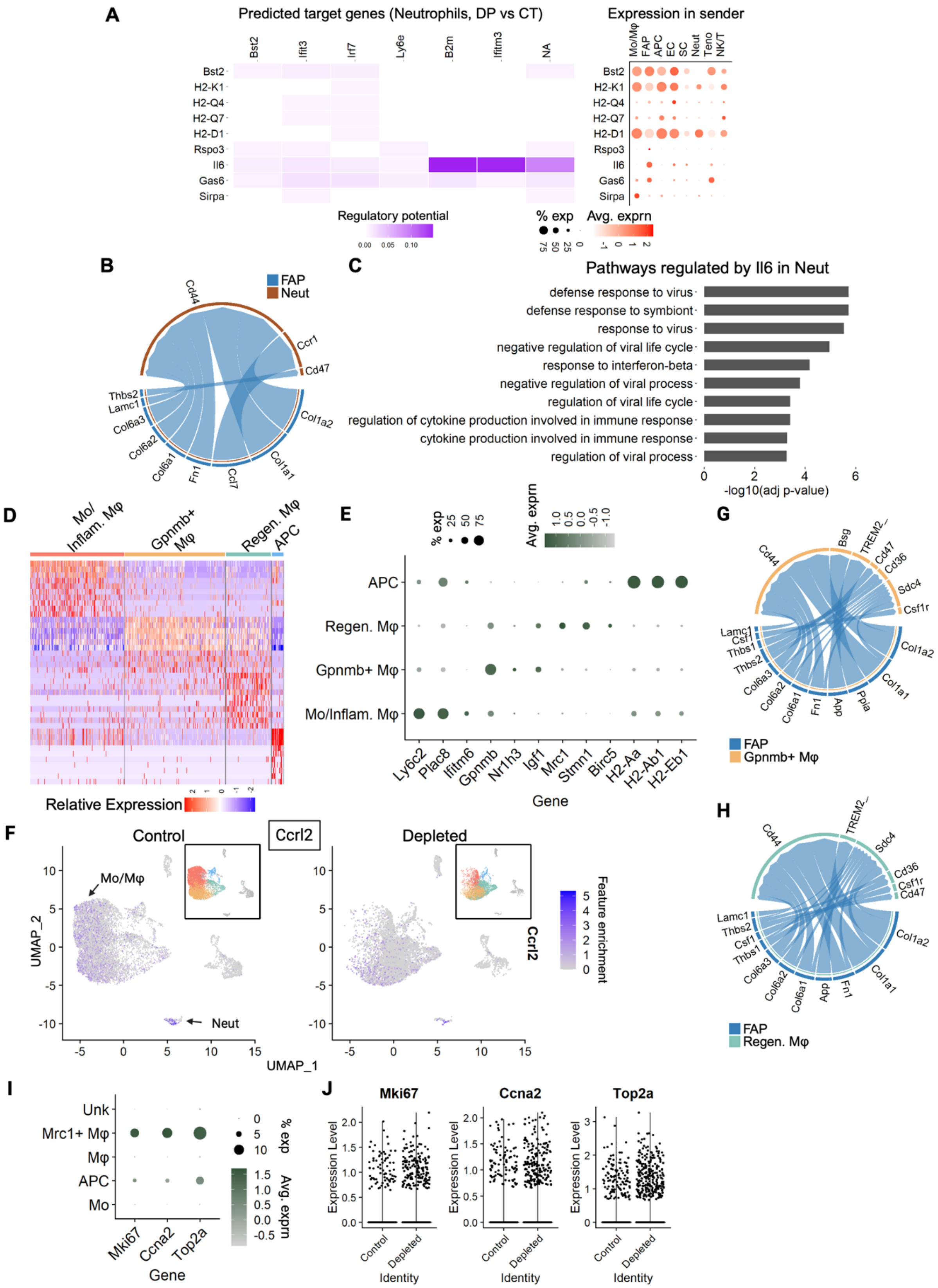
A
Predicted target genes (Neutrophils, DP vs CT)
Bst2
Ifit3
Irf7
Ly6e
B2m
Ifitm3
NA
H2-K1
H2-Q4
H2-Q7
H2-D1
Rspo3
Il6
Gas6
Sirpa
Regulatory potential
0.00 0.05 0.10
Expression in sender
Mo/Mφ
FAP
APC
EC
SC
Neut
Teno
NK/T
% exp
75 50 25 0
Avg. exprn
-1 0 1 2
B
FAP
Neut
Cd44
Ccr1
Cd47
Thbs2
Lamc1
Col6a3
Col6a2
Col6a1
Fn1
Ccl7
Col1a1
Col1a2
C
Pathways regulated by Il6 in Neut
defense response to virus
defense response to symbiont
response to virus
negative regulation of viral life cycle
response to interferon-beta
negative regulation of viral process
regulation of viral life cycle
regulation of cytokine production involved in immune response
cytokine production involved in immune response
regulation of viral process
0 2 4 6
-log10(adj p-value)
D
Mo/ Inflam. Mφ
Gpnmb+ Mφ
Regen. Mφ
APC
Relative Expression
2 1 0 -1 -2
E
% exp
25 50 75
Avg. exprn
1.0 0.5 0.0 -0.5 -1.0
APC
Regen. Mφ
Gpnmb+ Mφ
Mo/Inflam. Mφ
Ly6c2
Plac8
Ifitm6
Gpnmb
Nr1h3
Igf1
Mrc1
Stmn1
Birc5
H2-Aa
H2-Ab1
H2-Eb1
Gene
G
Cd44
Bsg
TREM2_
Cd47
Cd36
Sdc4
Csf1r
Col1a2
Col1a1
Ppia
App
Fn1
Col6a1
Col6a2
Col6a3
Thbs2
Thbs1
Csf1
Lamc1
FAP
Gpnmb+ Mφ
F
Control
Ccrl2
Depleted
Mo/Mφ
Neut
UMAP_2
UMAP_1
10 5 0 -5 -10
-5 0 5 10 15
Feature enrichment
5 4 3 2 1 0
Ccrl2
H
Cd44
TREM2_
Sdc4
Cd36
Csf1r
Cd47
Col1a2
Col1a1
Fn1
App
Col6a1
Col6a2
Col6a3
Thbs1
Csf1
Thbs2
Lamc1
FAP
Regen. Mφ
I
Unk
Mrc1+ Mφ
Mφ
APC
Mo
Mki67
Ccna2
Top2a
Gene
% exp
0 5 10
Avg. exprn
1.5 1.0 0.5 0.0 -0.5
J
Mki67
Ccna2
Top2a
Expression Level
Control
Depleted
Identity

**Supplementary Figure 12. FAP-dependent regulatory mechanisms of neutrophil and mononuclear phagocyte activity in SkM regeneration** (A) Top FAP-derived ligands with the highest activity and their predicted impact on DEGs in neutrophils, with the dot plot showing the abundance and expression level of these ligands across cell types in SkM at 3 dpi. (B) Interactions between FAPs (sender) and neutrophils (receiver). The thickness of lines represents interaction strength. (C) Top 10 biological processes in neutrophils (Neut) inferred to be regulated by Il6. (D) Heatmap showing the top 10 enriched features in each subpopulation of mononuclear phagocytes in regenerating SkM (3 dpi) resolved with the FindAllMarkers function. Specific markers are listed in Supplementary Table 3. (E) Dot plot representing the abundance and expression level of genes for subcluster identification across mononuclear phagocytes in SkM at 3 dpi. (F) UMAPs of cells in SkM at 3 dpi overlaid with the expression level of Ccrl2 in the noted samples. Inserts indicate the identities of mononuclear phagocytes by colors. Red - inflammatory Mo/Mφs; yellow - Gpnmb+ macrophages; green - regenerative macrophages; blue - APCs. (G, H) Interactions between FAPs (sender) and (G) Gpnmb+ macrophages and (H) regenerative macrophages (receivers). The thickness of lines represents interaction strength. The truncated term “TREM2_” = TREM2_TYROBP. (I) Dot plot representing the abundance and expression level of genes associated with proliferation in regenerative macrophages across cell types in SkM at 3 dpi. (J) Violin plots represent gene expression levels associated with proliferation in regenerative macrophages, split by control or FAP-depleted (Depleted) sample.

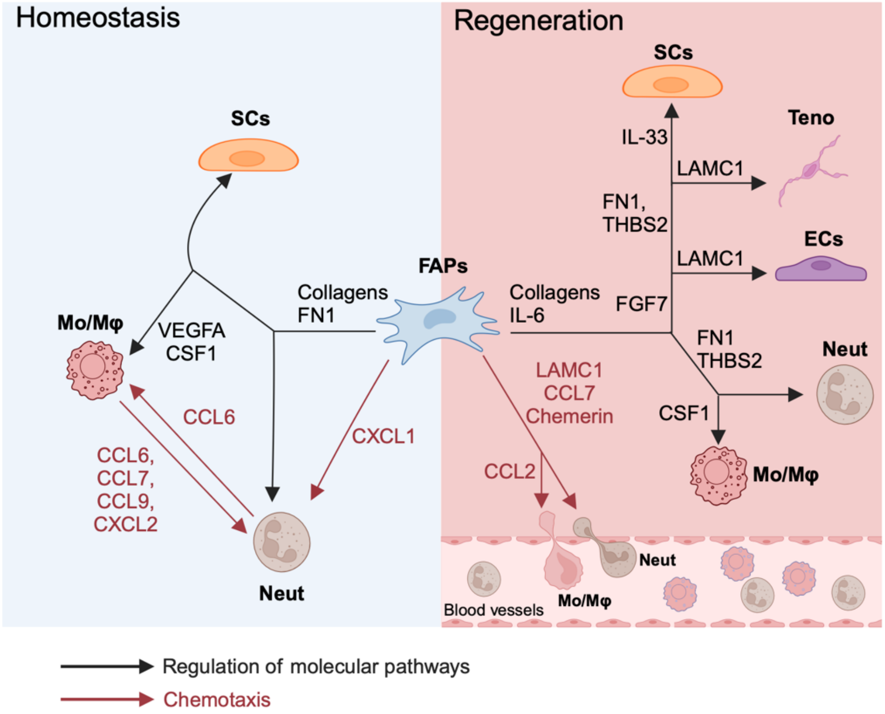


**Supplementary Figure 13. A FAP-centered interactome in homeostatic and regenerative SkM.**

**Supplementary tables**

**Table S1 (Gene List and Source Data for Figure S5B). Top10 Marker genes in each satellite cell subcluster in uninjured muscle.**

| cluster | gene | avg_log2FC | pct.1 | pct.2 | p_val | adj_pval |
|---|---|---|---|---|---|---|
| 0 | Ubb | 0.540749233 | 0.971 | 0.962 | 5.21E-19 | 5.49E-18 |
| 0 | Ubc | 0.61978907 | 0.95 | 0.92 | 3.32E-16 | 3.04E-15 |
| 0 | Hmox1 | 0.810980842 | 0.613 | 0.496 | 1.36E-07 | 4.74E-07 |
| 0 | Hspa1a | 0.477712133 | 0.704 | 0.588 | 1.31E-06 | 3.81E-06 |
| 0 | S100a10 | 0.507868038 | 0.789 | 0.776 | 4.40E-05 | 9.02E-05 |
| 0 | Arid5b | 0.375956789 | 0.872 | 0.849 | 0.000157713 | 0.000275443 |
| 0 | Txnrd1 | 0.653905726 | 0.505 | 0.462 | 0.000378709 | 0.000595544 |
| 0 | Crlf1 | 0.478196269 | 0.586 | 0.588 | 0.001624775 | 0.002062855 |
| 0 | Hspa1b | 0.480121289 | 0.435 | 0.368 | 0.002997447 | 0.003517509 |
| 0 | Nop53 | 0.356015497 | 0.681 | 0.7 | 0.005123171 | 0.005578009 |
| 1 | Mt2 | 1.170877865 | 0.983 | 0.895 | 5.04E-41 | 8.80E-40 |
| 1 | Mt1 | 1.267645095 | 0.963 | 0.797 | 4.80E-33 | 7.15E-32 |
| 1 | Ccn2 | 2.578481082 | 0.279 | 0.056 | 1.62E-23 | 1.90E-22 |
| 1 | Msc | 1.698013475 | 0.375 | 0.124 | 1.21E-18 | 1.25E-17 |
| 1 | Fas | 1.285985369 | 0.421 | 0.197 | 1.98E-13 | 1.44E-12 |
| 1 | Sod2 | 1.292703668 | 0.325 | 0.137 | 5.01E-11 | 2.88E-10 |
| 1 | Myf5 | 1.173624098 | 0.258 | 0.097 | 3.92E-10 | 2.02E-09 |
| 1 | Rhoh | 1.456511514 | 0.262 | 0.109 | 2.55E-09 | 1.16E-08 |
| 1 | Bgn | 1.299324672 | 0.358 | 0.181 | 5.12E-09 | 2.27E-08 |
| 1 | Igfbp5 | 1.333915892 | 0.396 | 0.221 | 2.92E-08 | 1.14E-07 |
| 2 | Col3a1 | 1.539834597 | 0.631 | 0.272 | 4.16E-24 | 4.99E-23 |
| 2 | Itm2a | 1.253665399 | 0.535 | 0.252 | 1.54E-15 | 1.33E-14 |
| 2 | Eif4ebp1 | 1.292652826 | 0.394 | 0.155 | 1.57E-13 | 1.16E-12 |
| 2 | Mthfd2 | 1.529965043 | 0.293 | 0.098 | 3.68E-13 | 2.60E-12 |
| 2 | Tagln | 1.681410625 | 0.288 | 0.094 | 5.74E-13 | 4.03E-12 |
| 2 | Cryab | 1.001463377 | 0.712 | 0.509 | 1.07E-10 | 5.82E-10 |
| 2 | Nars | 0.997734934 | 0.46 | 0.233 | 2.45E-10 | 1.30E-09 |
| 2 | Hsp90b1 | 0.937656777 | 0.596 | 0.357 | 6.56E-10 | 3.27E-09 |
| 2 | Mpp5 | 0.934510622 | 0.298 | 0.134 | 1.03E-07 | 3.70E-07 |
| 2 | Atf5 | 1.018052979 | 0.348 | 0.177 | 1.15E-07 | 4.09E-07 |
| 3 | Fos | 2.77179935 | 0.807 | 0.354 | 1.19E-32 | 1.75E-31 |
| 3 | Junb | 1.761292322 | 0.939 | 0.592 | 1.59E-31 | 2.22E-30 |
| 3 | Egr1 | 1.911231134 | 0.877 | 0.551 | 1.69E-26 | 2.22E-25 |
| 3 | Nr4a1 | 2.761407767 | 0.474 | 0.126 | 6.80E-25 | 8.43E-24 |
| 3 | Fosb | 1.682325288 | 0.789 | 0.497 | 3.09E-17 | 3.05E-16 |
| 3 | Gas1 | 2.226867733 | 0.491 | 0.181 | 4.76E-17 | 4.62E-16 |
| 3 | Socs3 | 2.115557134 | 0.491 | 0.189 | 1.04E-16 | 9.99E-16 |
| 3 | Hes1 | 2.592919802 | 0.307 | 0.117 | 1.17E-09 | 5.54E-09 |
| 3 | Gadd45b | 1.692080585 | 0.553 | 0.358 | 4.37E-09 | 1.94E-08 |

| | | | | | | |
|---|---|---|---|---|---|---|
| 3 | Tob1 | 1.706685798 | 0.289 | 0.122 | 2.48E-07 | 8.33E-07 |
| 4 | Slfn5 | 6.659967168 | 0.593 | 0.008 | 3.30E-112 | 1.84E-109 |
| 4 | Cd93 | 6.554090744 | 0.525 | 0.006 | 2.83E-102 | 4.50E-100 |
| 4 | Esam | 6.605175644 | 0.508 | 0.006 | 3.11E-98 | 3.48E-96 |
| 4 | Cyyr1 | 7.898091595 | 0.424 | 0.002 | 1.11E-91 | 1.12E-89 |
| 4 | Adgrl4 | 6.45356283 | 0.407 | 0.003 | 7.13E-84 | 4.68E-82 |
| 4 | Jam2 | 6.380089823 | 0.407 | 0.004 | 9.93E-81 | 5.54E-79 |
| 4 | Ablim3 | 6.582342521 | 0.339 | 0.003 | 2.29E-68 | 8.50E-67 |
| 4 | Mef2c | 7.437682265 | 0.322 | 0.003 | 1.31E-64 | 4.29E-63 |
| 4 | Itga6 | 6.590197601 | 0.322 | 0.004 | 1.62E-61 | 4.75E-60 |
| 4 | Acta1 | 7.870265357 | 0.254 | 0.039 | 6.41E-14 | 4.90E-13 |

**Table S2 (Gene List and Source Data for Figure S5F). Top10 Marker genes in each monocyte/macrophage subcluster in uninjured muscle.**

| cluster | gene | avg_log2FC | pct.1 | pct.2 | p_val | adj_pval |
|---|---|---|---|---|---|---|
| Infiltrating Mo | Hp | 4.00967197 | 0.87 | 0.066 | 4.95E-77 | 1.31E-73 |
| Infiltrating Mo | Sell | 4.150645 | 0.696 | 0.037 | 1.69E-63 | 1.12E-60 |
| Infiltrating Mo | Slpi | 4.75991369 | 0.783 | 0.086 | 2.38E-60 | 1.26E-57 |
| Infiltrating Mo | C3 | 4.63862533 | 0.574 | 0.04 | 6.18E-49 | 1.49E-46 |
| Infiltrating Mo | Mcemp1 | 3.88400691 | 0.617 | 0.059 | 1.98E-46 | 3.49E-44 |
| Infiltrating Mo | Prtn3 | 5.34717725 | 0.409 | 0.015 | 4.4E-38 | 5.3E-36 |
| Infiltrating Mo | Vcan | 4.23156547 | 0.374 | 0.033 | 1.5E-27 | 6.5E-26 |
| Infiltrating Mo | Mmp8 | 3.8107243 | 0.287 | 0.026 | 8.2E-21 | 1.5E-19 |
| Infiltrating Mo | Tarm1 | 3.94965627 | 0.278 | 0.024 | 1.7E-20 | 2.8E-19 |
| Infiltrating Mo | Plcb1 | 3.84685568 | 0.261 | 0.02 | 5.1E-20 | 8.1E-19 |
| IFNγ-sensitive Mφ | Cd74 | 1.84083803 | 0.905 | 0.634 | 1.02E-23 | 2.56E-22 |
| IFNγ-sensitive Mφ | H2-Eb1 | 1.81221745 | 0.77 | 0.372 | 2.47E-21 | 4.66E-20 |
| IFNγ-sensitive Mφ | Cxcl2 | 2.49885422 | 0.73 | 0.494 | 5.04E-15 | 4.99E-14 |
| IFNγ-sensitive Mφ | St3gal6 | 2.64069975 | 0.325 | 0.077 | 3.48E-14 | 3.22E-13 |
| IFNγ-sensitive Mφ | Pmp22 | 1.68495796 | 0.603 | 0.339 | 1.80E-12 | 1.30E-11 |
| IFNγ-sensitive Mφ | Tnfrsf12a | 1.76773804 | 0.429 | 0.257 | 3.89E-07 | 1.30E-06 |
| IFNγ-sensitive Mφ | Hpgds | 2.22940167 | 0.278 | 0.122 | 1.39E-06 | 4.10E-06 |
| IFNγ-sensitive Mφ | Ccl4 | 2.04606894 | 0.302 | 0.149 | 5.64E-06 | 1.47E-05 |
| IFNγ-sensitive Mφ | Ccl3 | 2.78866996 | 0.357 | 0.21 | 6.93E-06 | 1.78E-05 |
| IFNγ-sensitive Mφ | Creb5 | 1.7949592 | 0.286 | 0.14 | 1.23E-05 | 2.98E-05 |
| IFNβ-sensitive Mφ | Ifit3 | 4.80270221 | 0.583 | 0.023 | 1.12E-47 | 2.48E-45 |
| IFNβ-sensitive Mφ | Irf7 | 3.88756111 | 0.958 | 0.255 | 3.89E-35 | 3.68E-33 |
| IFNβ-sensitive Mφ | Ifi213 | 3.82043909 | 0.438 | 0.027 | 2.32E-29 | 1.18E-27 |
| IFNβ-sensitive Mφ | Ly6a | 3.71087137 | 0.771 | 0.134 | 4.04E-29 | 1.94E-27 |
| IFNβ-sensitive Mφ | Il18bp | 3.95170092 | 0.396 | 0.025 | 1.97E-26 | 7.43E-25 |
| IFNβ-sensitive Mφ | Slfn4 | 3.83129842 | 0.438 | 0.038 | 8.63E-25 | 2.66E-23 |
| IFNβ-sensitive Mφ | Ifi205 | 3.95175171 | 0.333 | 0.019 | 2.48E-23 | 5.90E-22 |
| IFNβ-sensitive Mφ | Cfb | 3.79259812 | 0.479 | 0.058 | 5.21E-23 | 1.18E-21 |
| IFNβ-sensitive Mφ | Arg1 | 4.06070313 | 0.375 | 0.036 | 3.31E-20 | 5.34E-19 |
| IFNβ-sensitive Mφ | Ifit1 | 3.82946078 | 0.292 | 0.031 | 8.48E-15 | 8.28E-14 |
| Tissue-resident Mφ | Lyve1 | 5.47420635 | 0.631 | 0.025 | 1.44E-59 | 6.33E-57 |
| Tissue-resident Mφ | Folr2 | 3.54262017 | 0.786 | 0.138 | 5.99E-45 | 9.91E-43 |
| Tissue-resident Mφ | C4b | 4.42717991 | 0.464 | 0.027 | 5.51E-38 | 6.18E-36 |
| Tissue-resident Mφ | Fxyd2 | 3.39924518 | 0.595 | 0.113 | 1.79E-29 | 9.28E-28 |
| Tissue-resident Mφ | Fcna | 4.51011926 | 0.31 | 0.014 | 6.56E-27 | 2.67E-25 |
| Tissue-resident Mφ | Fgfr1 | 3.50054193 | 0.417 | 0.049 | 4.47E-25 | 1.43E-23 |
| Tissue-resident Mφ | Cfh | 3.16740812 | 0.571 | 0.157 | 3.30E-22 | 7.05E-21 |
| Tissue-resident Mφ | Ccl8 | 3.67244107 | 0.369 | 0.045 | 1.86E-21 | 3.62E-20 |
| Tissue-resident Mφ | Cd163 | 3.48112948 | 0.405 | 0.087 | 7.81E-17 | 9.53E-16 |
| Tissue-resident Mφ | Clec10a | 3.7246707 | 0.345 | 0.07 | 3.25E-15 | 3.35E-14 |

| | | | | | | |
|---|---|---|---|---|---|---|
| Mφ | Mmp19 | 2.06266198 | 0.525 | 0.114 | 3.48E-24 | 9.80E-23 |
| Mφ | H2-DMb1 | 1.44362183 | 0.698 | 0.36 | 1.02E-15 | 1.14E-14 |
| Mφ | Il7r | 2.18935027 | 0.324 | 0.074 | 4.78E-14 | 4.35E-13 |
| Mφ | Flrt3 | 1.67694379 | 0.266 | 0.058 | 1.55E-11 | 1.01E-10 |
| Mφ | Fn1 | 1.64115834 | 0.633 | 0.372 | 2.68E-10 | 1.49E-09 |
| Mφ | Cd24a | 1.78399246 | 0.324 | 0.107 | 7.38E-10 | 3.88E-09 |
| Mφ | Cxcl3 | 1.55984943 | 0.324 | 0.109 | 3.99E-09 | 1.87E-08 |
| Mφ | Syngr1 | 1.52994331 | 0.273 | 0.081 | 4.00E-09 | 1.87E-08 |
| Mφ | Nr4a3 | 1.79630704 | 0.281 | 0.107 | 4.46E-07 | 1.48E-06 |
| Mφ | Irak1 | 1.57538215 | 0.266 | 0.156 | 8.11E-03 | 8.34E-03 |
| Unidentified | Cavin2 | 6.81160225 | 0.421 | 0.01 | 2.3E-41 | 3.6E-39 |
| Unidentified | Flt1 | 5.61045227 | 0.509 | 0.027 | 2.4E-40 | 3.2E-38 |
| Unidentified | Ace | 7.0454107 | 0.368 | 0.016 | 3.1E-31 | 1.9E-29 |
| Unidentified | Sparcl1 | 5.55895409 | 0.421 | 0.027 | 1.1E-30 | 6.7E-29 |
| Unidentified | C1qtnf9 | 5.60519111 | 0.316 | 0.01 | 6.7E-29 | 3.2E-27 |
| Unidentified | Gng11 | 4.99230007 | 0.439 | 0.041 | 2.5E-27 | 1E-25 |
| Unidentified | Col4a1 | 5.69123462 | 0.281 | 0.01 | 3.6E-25 | 1.2E-23 |
| Unidentified | Apold1 | 6.42178546 | 0.281 | 0.012 | 6.5E-24 | 1.7E-22 |
| Unidentified | Esam | 5.87883017 | 0.281 | 0.012 | 8.2E-24 | 2.1E-22 |
| Unidentified | Cldn5 | 5.79320866 | 0.263 | 0.014 | 7.5E-21 | 1.4E-19 |

Mo = Monocyte

Mφ = Macrophage

**Table S3 (Gene List and Source Data for Figure S12D). Top10 Marker genes in each mononuclear phagocyte subcluster in muscle at 3 day-post-injury.**

| cluster | gene | avg_log2FC | pct.1 | pct.2 | p_val | adj_pval |
|---|---|---|---|---|---|---|
| Mo/Inflam. Mφ | Ly6c2 | 3.883582449 | 0.84 | 0.283 | 0.00E+00 | 0.00E+00 |
| Mo/Inflam. Mφ | Plac8 | 3.355236097 | 0.841 | 0.349 | 0.00E+00 | 0.00E+00 |
| Mo/Inflam. Mφ | Ly6a | 2.181421004 | 0.689 | 0.344 | 0.00E+00 | 0.00E+00 |
| Mo/Inflam. Mφ | Isg20 | 2.441982103 | 0.469 | 0.171 | 0.00E+00 | 0.00E+00 |
| Mo/Inflam. Mφ | Clec4e | 2.601198542 | 0.458 | 0.165 | 0.00E+00 | 0.00E+00 |
| Mo/Inflam. Mφ | Slfn4 | 2.836710535 | 0.362 | 0.096 | 0 | 0 |
| Mo/Inflam. Mφ | Cfb | 2.238125233 | 0.422 | 0.16 | 0 | 0 |
| Mo/Inflam. Mφ | Ifitm6 | 3.379975045 | 0.295 | 0.049 | 0 | 0 |
| Mo/Inflam. Mφ | Slpi | 3.67482489 | 0.305 | 0.09 | 6.8785E-275 | 1.3918E-273 |
| Mo/Inflam. Mφ | F10 | 2.691191801 | 0.265 | 0.086 | 1.6843E-210 | 2.2388E-209 |
| Gpnmb+ Mφ | Syngr1 | 1.848139692 | 0.895 | 0.371 | 0.00E+00 | 0.00E+00 |
| Gpnmb+ Mφ | Gpnmb | 1.31348491 | 0.865 | 0.421 | 0.00E+00 | 0.00E+00 |
| Gpnmb+ Mφ | Trem2 | 1.395006193 | 0.976 | 0.573 | 0.00E+00 | 0.00E+00 |
| Gpnmb+ Mφ | Timp2 | 1.142652717 | 0.919 | 0.583 | 0.00E+00 | 0.00E+00 |
| Gpnmb+ Mφ | Spp1 | 1.328859289 | 0.939 | 0.663 | 0.00E+00 | 0.00E+00 |
| Gpnmb+ Mφ | Ctsd | 1.20958679 | 0.997 | 0.859 | 0.00E+00 | 0.00E+00 |
| Gpnmb+ Mφ | Igf1 | 1.426537187 | 0.417 | 0.159 | 1.67E-255 | 2.98E-254 |
| Gpnmb+ Mφ | Abca1 | 1.162669772 | 0.462 | 0.188 | 5.55E-248 | 9.36E-247 |
| Gpnmb+ Mφ | Itgb5 | 1.15996018 | 0.475 | 0.207 | 5.69E-232 | 8.17E-231 |
| Gpnmb+ Mφ | Slc6a8 | 1.201644582 | 0.277 | 0.11 | 7.73E-141 | 5.94E-140 |
| Regen. Mφ | Stmn1 | 3.192374308 | 0.535 | 0.101 | 0.00E+00 | 0.00E+00 |
| Regen. Mφ | Igfbp4 | 3.104721663 | 0.436 | 0.083 | 0.00E+00 | 0.00E+00 |
| Regen. Mφ | Mrc1 | 2.473195519 | 0.464 | 0.125 | 0.00E+00 | 0.00E+00 |
| Regen. Mφ | Tubb5 | 1.756694589 | 0.726 | 0.401 | 0.00E+00 | 0.00E+00 |
| Regen. Mφ | Birc5 | 4.494357402 | 0.265 | 0.017 | 0.00E+00 | 0.00E+00 |
| Regen. Mφ | Cdca3 | 4.184124402 | 0.25 | 0.018 | 0.00E+00 | 0.00E+00 |
| Regen. Mφ | Cbr2 | 2.115383851 | 0.473 | 0.16 | 8.00E-296 | 1.87E-294 |
| Regen. Mφ | Cks1b | 2.625085134 | 0.282 | 0.056 | 8.62E-279 | 1.82E-277 |
| Regen. Mφ | F13a1 | 1.859543271 | 0.494 | 0.196 | 1.84E-242 | 2.93E-241 |
| Regen. Mφ | Ccdc34 | 2.04285572 | 0.295 | 0.088 | 4.56E-186 | 5.12E-185 |
| APC | H2-Eb1 | 4.645908121 | 0.969 | 0.219 | 0.00E+00 | 0.00E+00 |
| APC | H2-Aa | 4.324019288 | 0.975 | 0.267 | 0.00E+00 | 0.00E+00 |
| APC | H2-Ab1 | 4.363090025 | 0.969 | 0.283 | 0.00E+00 | 0.00E+00 |
| APC | Gpr171 | 6.012723757 | 0.369 | 0.01 | 0.00E+00 | 0.00E+00 |
| APC | Klrd1 | 9.064173594 | 0.329 | 0.001 | 0.00E+00 | 0.00E+00 |
| APC | Ccnd2 | 4.098551847 | 0.335 | 0.028 | 0.00E+00 | 0.00E+00 |
| APC | Ramp3 | 5.702406079 | 0.311 | 0.008 | 0.00E+00 | 0.00E+00 |
| APC | H2-DMb2 | 5.763229858 | 0.3 | 0.01 | 0.00E+00 | 0.00E+00 |
| APC | Ccr7 | 9.734511728 | 0.262 | 0.002 | 0.00E+00 | 0.00E+00 |

* In R, any number smaller than approximately 2.225074e-308 will be displayed as 0 due to floating-point limitations.

Mo = Monocyte

Mφ = Macrophage

Inflam. = Inflammatory

Regen. = Regeneration
